# GPR17 structure and agonism with small molecules and oxysterols

**DOI:** 10.1101/2025.11.26.690757

**Authors:** Claire M. Metrick, Lu Zhang, Maria Hercher, Gregory J. Dodge, Dennis Cherry, Jorge Vera Rebollar, Margot Brickelmaier, Jing Wang, Jeffrey G. Martin, Xiaoping Hronowski, Melissa Kemp, Andrew Capacci, Joseph W. Arndt, Laura Silvian, Karen S. Conrad, John Silbereis

## Abstract

GPR17 is a class A orphan GPCR that regulates myelination in the central nervous system. Inhibiting GPR17 is a potential therapeutic mechanism to induce remyelination. The limited understanding of the structural basis of GPR17 agonism and antagonism is a challenge for GPR17 drug discovery. We present novel cryo-EM structures of GPR17 with bound orthosteric antagonist or agonist/G protein heterotrimer with biophysical and cellular characterization to inform rational drug discovery. Both ligands bind towards a lateral edge of the orthosteric pocket, while an exceptionally long extracellular loop 2 occupies and limits access to this site. In contrast, pharmacological and photoaffinity studies indicate that endogenous oxysterol agonists of GPR17 bind allosterically. Characterization of oxysterol agonism versus a known orthosteric ligand in oligodendrocytes further supports an allosteric mechanism. Therefore, the design of more potent orthosteric ligands is limited by pocket size and accessibility. Targeting allosteric binding offers an alternative approach to GPR17 antagonism.

## Introduction

GPR17 is an orphan G-protein coupled receptor (GPCR) that is predominantly expressed in the central nervous system (CNS)^1^. In the CNS, GPR17 is expressed in oligodendrocyte progenitor cells (OPCs), which mature into myelinating oligodendrocytes.^1–3^ The primary role of GPR17 is hypothesized to prevent terminal oligodendrocyte maturation to appropriately time myelination during development.^1^ In addition, GPR17 in cells is rapidly upregulated in response to demyelination in MS lesions, brain injuries, and ischemic infarct.^4–6^ Expression of GPR17 in demyelinating lesions has been shown to block efficient remyelination.^1,2,7–9^ Owing to these properties, GPR17 inhibition is a widely pursued drug target for remyelinating therapy.^9^

GPR17 is homologous to CysLT and P2Y receptors, whose native ligands include cysteinyl leukotrienes and nucleotides, and EBI2, an oxysterol sensor.^10^ Due to the homology to the CysLT and P2Y receptors, it was originally thought that their endogenous ligands could activate GPR17, but these observations failed to replicate in subsequent studies.^3,11,12^ Recent characterization of oxysterol binding and induced activity identified 24S-hydroxysterol as a proposed native ligand of GPR17, in line with the EBI2 mechanism of action.^13^ However, this too remains to be replicated in subsequent studies and native cell types. Thus, GPR17 remains an orphan GPCR despite several investigations of ligand binding. The consensus synthetic agonist MDL29,951 (2-carboxy-4,6-dichloro-1*H*-indole-3-propionic acid), has been shown to activate GPR17, engage G proteins via Gα_i_, Gα_s_ and Gα_q_ pathways^11^, and has been used extensively to study the biology and pharmacology of GPR17. Notably, MDL29,951 (MDL) selectively activates GPR17 in primary oligodendrocytes, thereby inhibiting the maturation of primary oligodendrocytes and model animal systems. This effect is abrogated by GPR17 knockout.^11^ The recent solution of the apo structure of GPR17 identified two features that enable constitutive activity including the ECL2 occupation of the orthosteric binding site and the altered sodium ion site. ^14^ However, constitutive activity may also be attributed to a naturally abundant native ligand like 24S- hydroxysterol which is found in rat and pig brain at pharmacologically-relevant levels,^13^ with additional quantitation in mouse and human in this work for validation.^15^ In the apo-structure of activated GPR17^16^, the ECL2 is anchored deep in the orthosteric pocket, making the docking of a large hydrophobic oxysterol in that pocket, like those seen in known sterol receptors SMO, GPR183, or EBI2, questionable despite the attempts to do so in earlier published GPR17 homology models and supportive pharmacological evidence. In sum, there are significant gaps in the understanding of the mechanistic and structural mechanism of GPR17 agonism, and a biophysical basis for GPR17 antagonism is yet to be determined.

Here we present structures of agonist (MDL)-bound and antagonist (I-185, 6-chloro-N-(4-cyano-2,5- difluoro-phenyl)-1H-indole-3-sulfonamide)-bound GPR17^17^, to clarify ligand binding mode and establish limitations of mobility of the ECL2-occupied orthosteric pocket. We determine that I-185 and MDL directly compete at the GPR17 orthosteric binding site and we identify sterol-like densities (SLD) flanking the transmembrane helices in the experimental map. In combination with photoprobe-based competition and mass spectrometry binding assays, we hypothesize that these SLDs represent allosteric binding sites for 24S-OHC or other sterols. Schild analysis reveals distinct pharmacological properties of GPR17 antagonism to MDL and oxysterol ligands that are respectively consistent with orthostery versus allostery. We then confirm activity of these ligands in native GPR17-expressing cell types: primary mouse oligodendrocytes and human IPSC-derived oligodendrocytes. Finally, we measure the levels of GPR17 and 24S-hydroxycholesterol in human control and multiple sclerosis brain samples to confirm that 24S-OHC may be present to agonize the receptor at biologically relevant concentrations.

## Results

### GPR17 structure determination

#### Cryo-EM structure of active GPR17 bound to G_i_ heterotrimer and small molecule agonist

The structure of GPR17 was solved in its active state in complex with Gα_i1_/β_1_/γ_2_ in the presence of agonist MDL to understand the modulation of its architecture from constitutively active to fully active state. We determined the structure of the complex at 3.5 Å resolution by cryo-EM (Fig. 1; Extended Data Fig. 1a, 2a,c,e). In the MDL-bound form, differences from the apo-activated-state structure are present that may indicate that the agonist-bound structure is in a fully activated form in comparison to its reported “self- activated” state. It is important to note that the density of the extracellular side of the receptor is more resolved in this MDL-bound structure than that of the published apo structure, likely due to the spatial restrictions from the ligand. The overall RMSD with the published apo-structure including G-proteins is 0.9 Å. The apparent structural changes are primarily around the ligand binding pocket, closest to the extracellular side, with minimal changes to the intracellular side and interaction with Gα_i_. Specifically, residues H220^5.38^ and S218^5.36^ move to make hydrogen bonds with one carboxylic acid moiety of MDL, and R283^6.55^ shifts to interact with the second carboxylic acid, stabilized by a new contact with Y279^6.51^(Fig. 1a). The phenyl ring of MDL has a sulfur-aromatic interaction with M191^4.57^ in addition to both a pi-pi interaction with Y144^3.37^ and an interaction between the hydroxyl group of Y144^3.37^ and one MDL chlorine. Finally, S224^5.42^ shifts to interact with the indole nitrogen. These interactions provide a tightened region surrounding the ligand, pulling the extracellular ends of all transmembrane helices (TMs) and the second extracellular loop (ECL2) toward the ligand, with slight rotation in TM4 and TM5.

**Figure 1.**
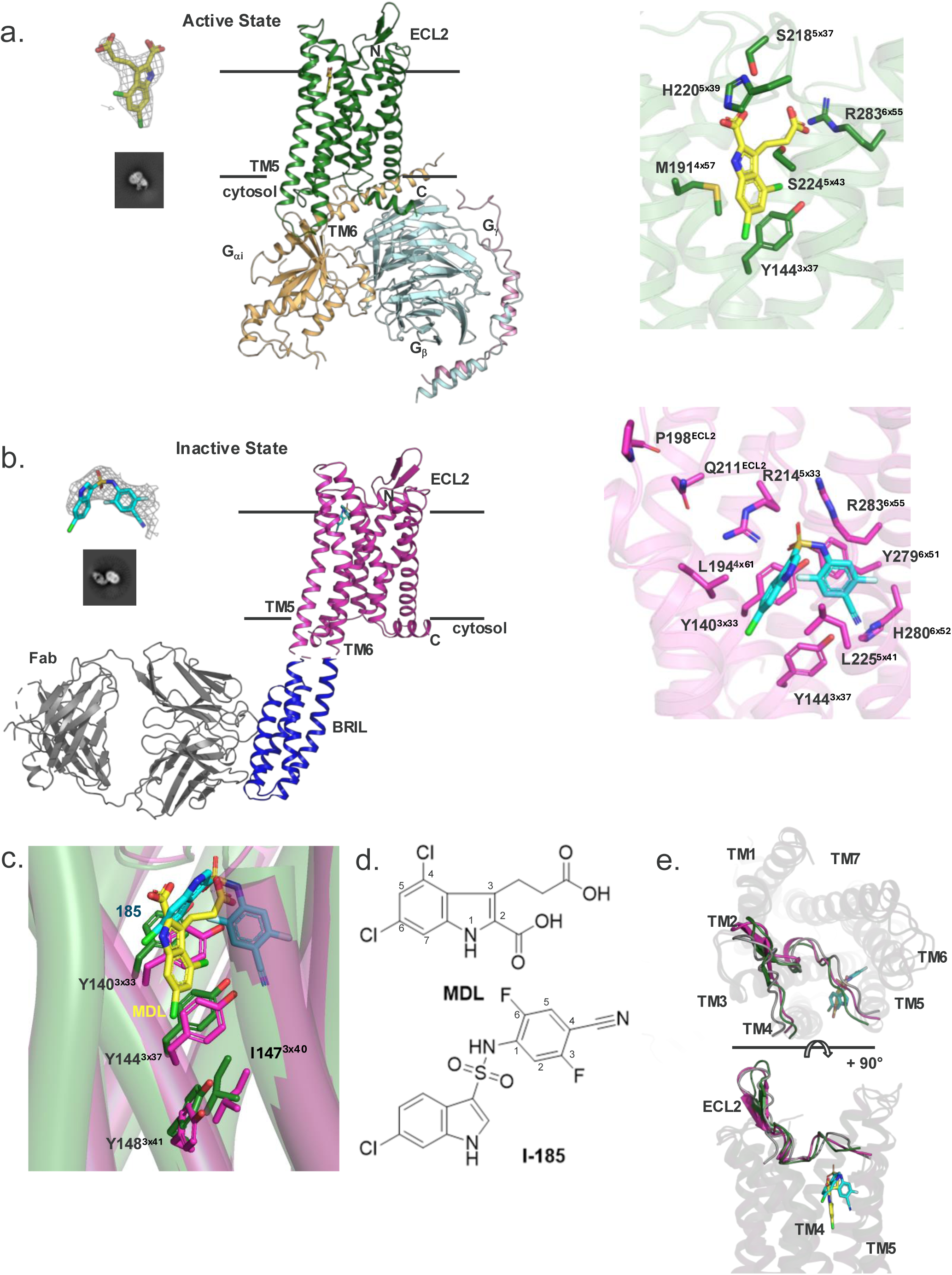
Structure Determination of Active and Inactive GPR17. a) Active state structure of GPR17 in complex with Gαi1/β1/γ2 and MDL. Also shown: MDL (yellow stick) in cryoEM density, representative 2D class, and proposed binding mode. b) Inactive state structure of bRIL-modified GPR17 in complex I-185. Also shown: I-185 (cyan stick) in cryoEM density, representative 2D class, and proposed binding mode. c) Binding modes of MDL (yellow sticks in green ribbon, left) and I-185 (cyan sticks in fuchsia ribbon, center) overlaid, with residues impacting microswitch cascade (cylinders, right). d) Molecular structures of MDL and I-185 with atoms numbered. e) ECL2 from apo (grey, PDB: 7Y89), MDL-bound (green), and I-185-bound (fuchsia) GPR17 overlaid.

Based on the apo structure, it was suggested that the ECL2 fills part of the orthosteric pocket resulting in constitutive activity. Our structure shows that in a small molecule agonist-activated GPR17, ECL2 remains in place, occupying part of the orthosteric pocket with the remainder filled by MDL. The interactions of the ECL2 with the orthosteric pocket identified in the apo structure are in place, including its hydrogen bonds with TM7 and disulfide with TM3. The improvements in the map of the MDL-bound structure clarify the conformations of the loop and hairpin in the extracellular region. Our structure with MDL reinforces the ECL2 location in the orthosteric pocket from the apo structure and adds the observation that this position persists in the presence of small molecule agonist (Fig. 1e). Without knowledge of the unmoving ECL2, the binding modes of previously identified GPR17 agonists^18^ were illustrated structurally using a homology model from related GPCR P2Y_12._^19^ Each chemotype was placed in a unique, but more central position among the 7 TMs consistent with the established class A GPCR orthosteric site. With our MDL-bound experimental structure, we define one common binding mode to rationalize the reported SAR.

The experimental position of MDL shows us that the 6- position provides a vector leading into a hydrophobic valley comprising Y148^3.41^, L183^4.49^, V186^4.52^, V187^4.53^, A228^5.46^, and P232^5.50^ that is accessible to the membrane between TM4 and TM5. Antagonist-bound structures of CysLT2^20^ and CysLT1^21^ show long lipophilic chains extending from the orthosteric ligands into this cleft. Members of the δ-subgroup of GPCRs, including EBI2 (GPR183)^22^ have a putative hydrophobic substrate entrance in this cavity that would equate to P193^4.59^, A190^4.55^ and S224^5.42^ (Extended Data Fig. 3).^22^

#### CryoEM structure of small molecule antagonist-bound inactive GPR17

The structure of GPR17 was also solved in its inactive state by cryo-EM to 3.32 Å resolution using a negative modulator I-185 and a modified construct with truncated N- and C-termini, a thermostabilized apocytochrome b_562_RIL^23^ insertion fused with A2a-based rigid linkages between TM5 and TM6^24^, and two thermostabilizing mutations, D105^2.50^N and F158^3.51^Y,^25^ in complex with an anti-bRIL Fab^26^ and anti-Fab nanobody (Fig. 1, Extended Data Fig. 1b, 2b,d,f). These stabilizing mutations were designed from referencing the mutations used to thermostabilize the CysLT2 receptor for antagonist-bound structural characterization.^20^ This enabled a unique structural characterization of an inactivated, constitutively active GPCR.

Our I-185 bound structure shows that the ECL2 remains in the orthosteric pocket during inhibition, with I-185 filling a similar space to MDL (Fig. 1c). In the antagonist-bound form, the I-185 ligand is bound through both electrostatic and hydrophobic interactions within the orthosteric binding pocket, and the protein is in a consensus inactive state including a large shift inward of the intracellular side of TM6 (Fig. 2a). In this placement there are polar contacts from R283^6.55^ and the ECL2 R214^5.32^ to the sulfonamide that anchors this antagonist (Fig. 2c). R214^5.32^ also interacts with Q211^ECL2^, which is in turn stabilized by the backbone at P198^4.59^. The nitrile moiety likely interacts with H280^6.52^ and potentially Y144^3.37^, the phenyl ring is sandwiched between L225^5.40^ and a hydrophobic patch comprising Y140^3.33^, Y279^6.51^, and the aryl arm of R283^6.55^. Similarly, the indole ring is aligned between L225^5.40^ and L194^4.60^ but also contacts S218^5.36^ through the indole nitrogen. Notably, the indole moieties of MDL and I-185 do not overlap but are shifted in position (Fig. 2c). The position of I-185 also allows for a second potential access point between helices 5 and 6 due to a bending of the extracellular side of TM5 towards TM6. Unlike its related δ-GPCR P2Y_12_, there are no radical changes from active to inactive states in the ECL2 conformation (Extended Data Fig. 5). Instead, similar to EBI2, the ECL2 fills the orthosteric cavity of GPR17 similarly in the active and inactive conformations. There are also similarities in the ligand placement of antagonist-bound CysLT2/1 structures (Extended data Fig. 3). Like for MDL-related agonists, SAR observations from various GPR17 antagonist scaffolds have been described and illustrated through docking into the orthosteric site of P2Y_12_-based homology models of GPR17.^27,28^ Again, these models do not account for the occupation of the orthosteric pocket with the ECL2, and specific proposed interactions do not appear to be valid. However, the general SAR observations may still be feasible when their molecules are placed in the binding site of I-185.

**Figure 2.**
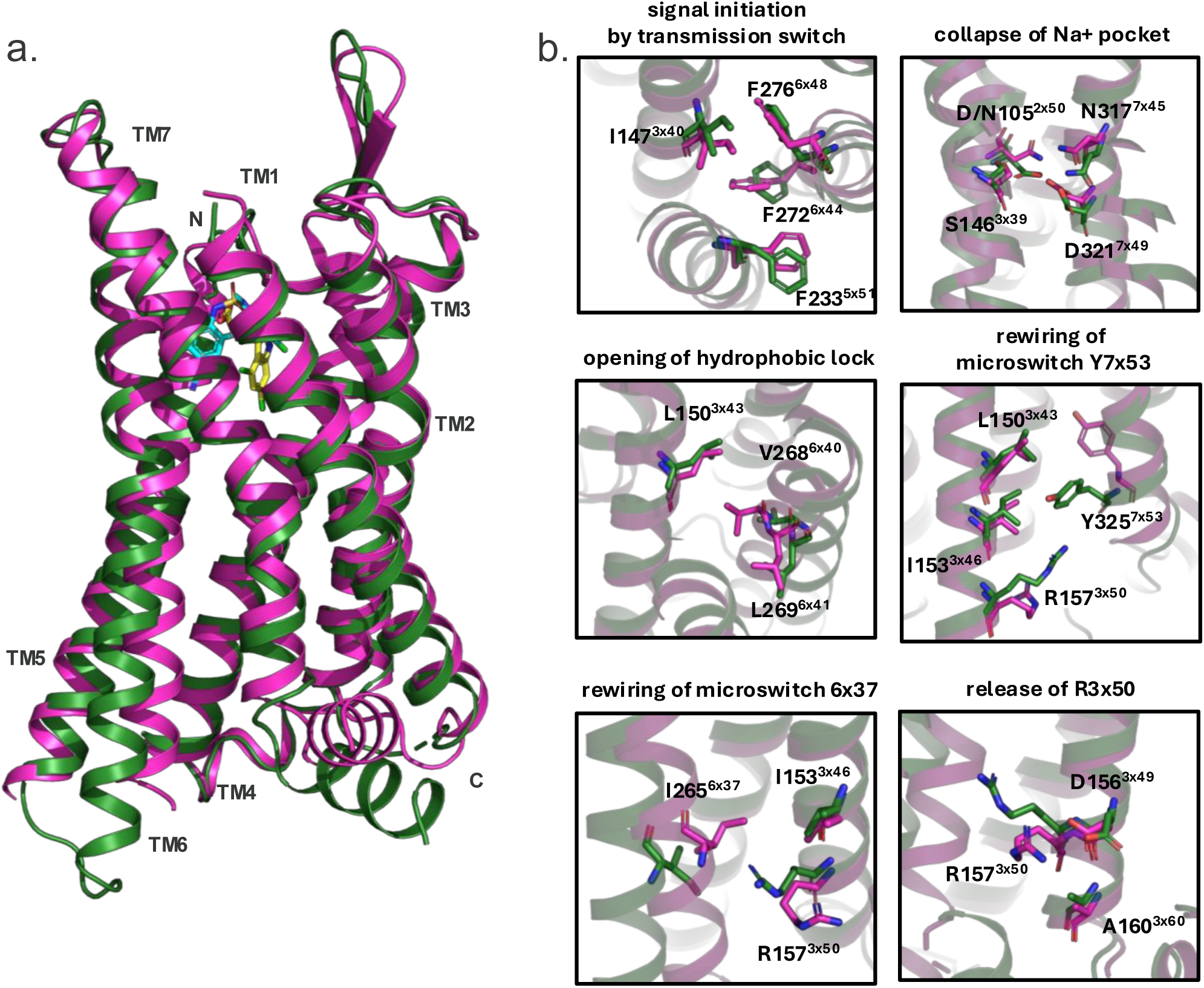
Molecular Mechanism of GPR17 (In)activation. a) Ribbon diagrams of MDL-bound (green) and I-185-bound (fuchsia) GPR17 overlaid. b) Comparison of known microswitch residues between MDL-bound (green) and I-185-bound (fuchsia) GPR17.

A comparison of the antagonist- and agonist-bound structures including microswitches involved in the consensus activation mechanism of class A GPCRs demonstrates the distinct states of GPR17 activation (Fig. 2b). Activation signal is initiated by changes in the transmission switch with significant shifts in I147^3.40^, F276^6.48^ and F272^6.44^ as well as the collapse of the Na^+^ pocket D/N105^2.50^, S146^3.39^, N317^7.45^ and D321^7.49^. There is rewiring of Y325^7.53^, I265^6.37^ and R157^3.50^ (DRY motif). The stabilizing mutations in the inactive state structure D105N^2.50^ and F158Y^3.51^ respectively remove charge that stabilizes the Na^+^ pocket activation switch^29^ and return the DRF motif to the more conserved DRY ionic lock of class A GPCRs.

#### ECL2 orientation does not change in small molecule activated and inactivated states

The role of ECL2 in self-activation of GPR17 was validated in a previous report^16^ by analysis of ECL2 mutations. Either point mutations or multiple residue linker replacement significantly reduced activity compared to wildtype (WT) without significantly altering the surface expression. In GPR17, the ECL2 loop is located deep in the orthosteric pocket mediated by hydrogen bonding between Y213^ECL2^ and R308^7.36^ and R214^ECL2^ and N307^7.35^ in the apo structure. The R214 ^ECL2^ interaction as well as the interactions between residues in the ECL2 beta hairpin (V201-V207 ^ECL2^, Q202-T206 ^ECL2^, N204-T206 ^ECL2^), and C132^3.25^-C209^ECL2^ disulfide bond are conserved with ligand-bound structures. The R308^7.36^ interaction with Y213^ECL2^ subsides, as its side chain rotates, however Y213 ^ECL2^ does not move significantly on account of hydrogen bonding with H119^2.64^. The ECL2 is remarkably stable in its position in the orthosteric pocket and the conformational changes that accompany activation state are driven by the placement of the indole in the remaining portion of the pocket. Specifically, the indole impacts the position of Y140^3.33^ and Y144^3.37^ on TM3, a helix with ties to each microswitch, with side chain shifts that propagate down the helix to Y148^3.41^ and neighboring residue I147^3.40^, triggering the transmission switch and the rest of the canonical microswitch cascade excluding the sodium site (Fig. 2b-c). With the exception of the microswitches and TM6 movements that characterize class A GPCR inactivation, the transition from activated and constitutively active (apo) to inactivated states of GPR17 are rather subtle, with RMSD values between the 7TM domains under 1.5 Å (apo:MDL = 1.05 Å; apo:I-185 = 1.37 Å; and MDL:I-185 = 1.35 Å) and the ECL2 with RMSD values under 2.5 Å (apo:MDL = 2.05 Å; apo:I-185 = 1.29 Å; and MDL:I-185 = 1.58 Å). This stability is in stark contrast to the conformational changes that occur upon inactivation of many GPCRs, including other homologous receptors like P2Y_12_ (Extended Data Fig. 5). This is consistent, however, with the rigidity of the ECL2s of other self-activating GPCRs including GPR52 and GPR161. Though the specific interactions of ECL2 with the orthosteric pocket vary, the ECL2 does not make significant shifts with or without compound bound^14,30^ in GPR52 or in GPR161 MD simulations with or without miniG_s_ bound.^31^ It was shown by the addition of the Gα_s_ protein in the GPR52 cryoEM structure with C17 ligand, that the ligand binding pocket increases in size primarily by rotation of side chains^32^ as opposed to ECL2 movement.

### In vitro pharmacology

#### Functional Characterization of MDL-29951 and I-185 Interactions

Having established a structural basis for GPR17 antagonism at its orthosteric site, we next characterized the pharmacology of MDL and I-185. We generated a 1321N1 astrocytoma cell line stably expressing GPR17. Using this cell line, we determined the activity and potency of MDL using a homogenous time resolved fluorescence (HTRF)-based assay for cAMP detection in response to G_i_ coupled GPCRs (Fig. 3a,b). No activity was observed in wild type parent cells without GPR17 expression (data not shown). To define the nature of competition between MDL and I-185, we conducted a Schild analysis in which I-185 antagonist dose responses (potency and E_max_) were determined against increasing concentrations of MDL agonist. I-185 demonstrated a parallel right shift of dose response curves with increasing concentrations of MDL, but there was little impact on E_max_ (Fig. 3c,d). Schild regression analysis for ligand binding of MDL and I- 185 confirmed a linear relationship with a slope of 1.17 (Fig. 3d). Taken together, these results are consistent with competitive binding.^33^ Thus, we provide functional confirmation of direct competition of MDL and I-185 at the orthosteric site which matches the overlap of the agonist and antagonist in the overlayed structures (Fig. 1c).

**Figure 3.**
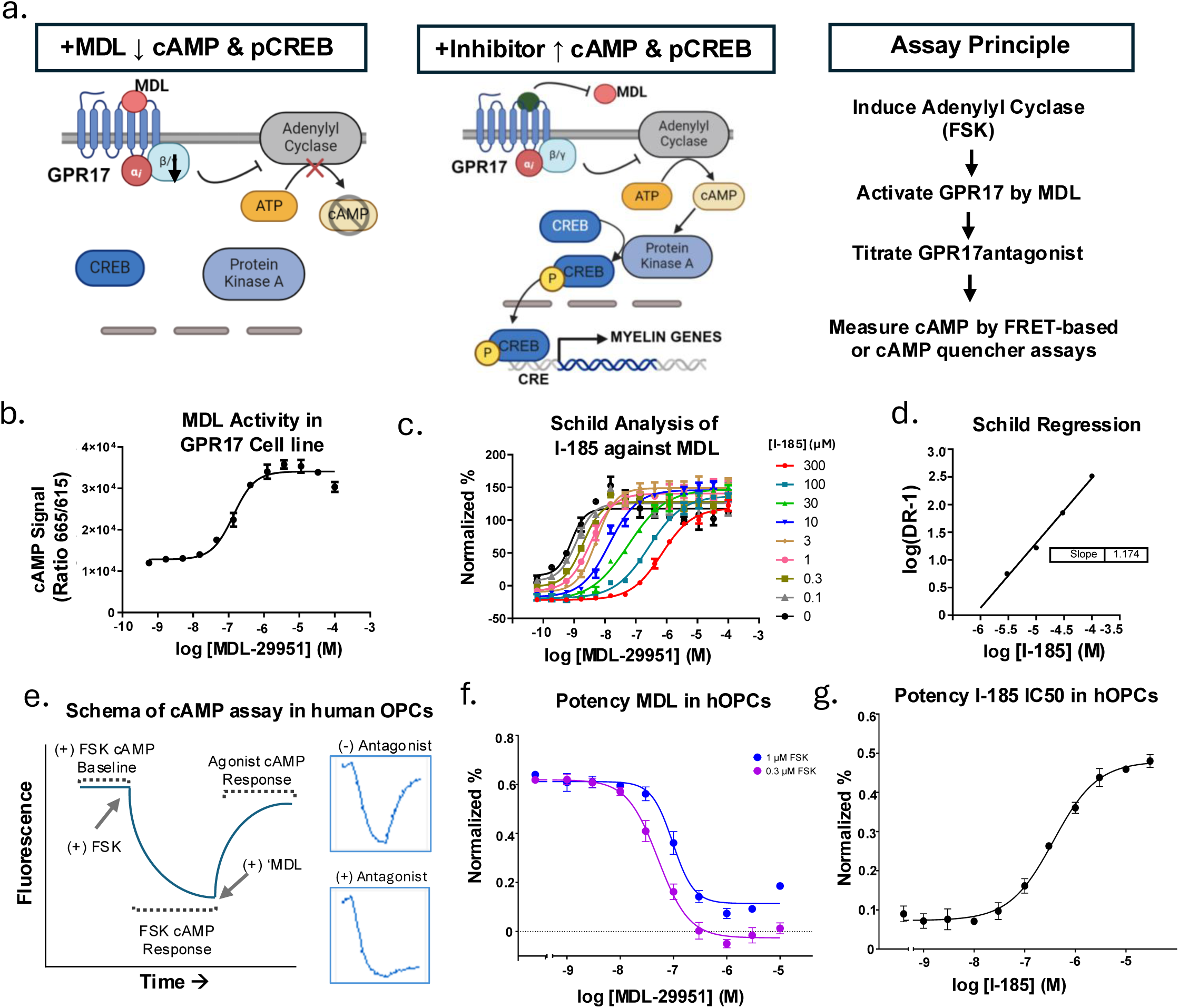
Pharmacology of I-185 and MDL-29951 interactions. a) Diagram of GPR17 signaling and assay principle for measuring cAMP as readout for GPR17 activity. b) Potency of MDL was determined in a 1321N1 astrocytoma line stably expressing GPR17. Normalized percentage is the change in signal from agonist relative to forskolin alone. c) Dose response curves in Schild analysis of I-185 versus MDL showing a right shift in parallel pattern of I-185 with increasing concentrations of MDL. d) Schild regression showing a linear agonist versus antagonist relationship indicative of competitive binding. e) Schematic of a BacMam-based cAMP sensor assay to measure I-185 and MDL activity in human OPCs. f) EC50 determination of MDL in hOPCs. Normalized data are expressed as the percentage of fluorescent signal relative to the baseline forskolin signal pre-agonist addition. g) IC50 determination of I-185 against MDL in hOPCs. Normalized data are expressed as the percentage ratio of fluorescent signal after antagonist addition divided by the difference in FSK baseline signal to agonist signal. Plots and error bars represent the mean ± SEM. *N* = 4 independent experiments with 2-4 technical replicates. Schematics were created in BioRender.

To extend the biological relevancy of MDL and I-185 ligands, we next characterized their activity in io.OPC human oligodendrocyte cultures (the native GPR17 expressing cell type). These cells are iPSC- derived human OPCs (hOPCs) with an engineered Tet-inducible transcription factor program for oligodendrocyte differentiation. We performed a differentiation time course and determined that GPR17 expression peaked at 5 days in culture and showed a gene and protein expression profile consistent with preOLs (Extended Data Fig. 7). Therefore, experiments were performed at day 5 in culture.

We found that these hOPC cultures were not amenable to miniaturization into the HTRF cAMP assay format. Therefore, we adapted a vector-based assay in which a BacMam virus was used to deliver a G_i_- coupled cAMP sensor system to the hOPCs (Fig. 3e). We determined that MDL has a similar potency and efficacy in hOPCs as in the GPR17 cell line (Fig. 3f). Antagonism of MDL by I-185 was comparable to that of the GPR17 cell line (Fig. 3c and g), confirming the MDL and I-185 interactions extend to the relevant cell lineage.

#### Effects of I-185 and MDL Interactions on GPR17 Signaling in OPCs

Functionally, G_i_-coupled GPCRs act mainly by inhibiting pCREB-mediated gene expression downstream of cAMP-PKA signaling (Fig. 4a). In oligodendrocyte development, pCREB signaling is a key activator of the transcriptional program for myelination and has been proposed as the possible mechanistic basis for inhibition of myelination by GPR17 activation^2,7,34^.

**Figure 4.**
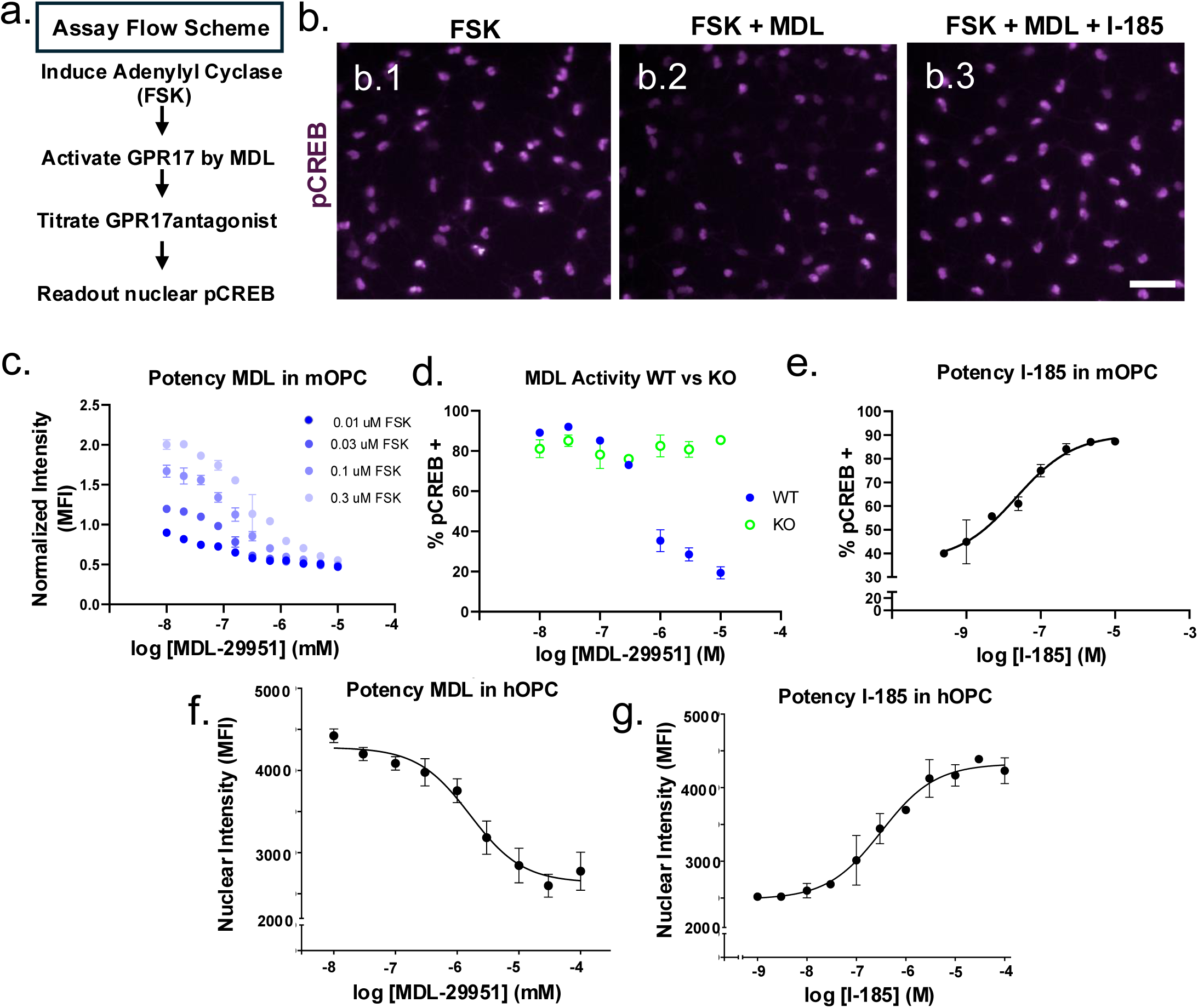
Effects of I-185 and MDL Interactions on GPR17 Signaling in OPCs. a) Assay principle and flow scheme for measurement of nuclear pCREB as a readout of GPR17 signaling. b) Representative immunohistochemical images demonstrating induction of nuclear pCREB downstream of GPR17 by Forskolin (FSK) activation of adenylyl cyclase (b.1); reduction of FSK-induced pCREB signal by MDL-29951 (b.2); and antagonism of MDL-29951 effect by I-185 (b.3). c) Determination of MDL potency against increasing levels of FSK. d) MDL activity in GPR17 wild type and knockout derived mouse OPCs. e) Dose response curve of I-185 against MDL. f) IC50 of MDL in human OPCs. g) I-185 dose response in human OPCs. Plots and error bars represent the mean ± SEM. *N* = 4 independent experiments with 2 to 4 technical replicates. Scale bar in b.3 = 50 µm.

To determine the effects of MDL and I-185 interaction on pCREB signaling downstream of GPR17, we developed a high content imaging assay for CREB phosphorylation and nuclear localization in mouse and human oligodendrocytes (Fig. 4). The nuclear localization of pCREB in response to GPR17 agonism by MDL was first assessed in mouse oligodendrocytes, where GPR17KO-derived OPCs allowed us to determine the specificity of the assay. FSK was used to induce nuclear pCREB. We then confirmed the ability of MDL to reverse FSK induction of pCREB in wild type by not KO mouse OPCs (Fig. 4 b-d; Extended Data Fig. 8). Dose response studies of MDL and I-185 confirmed I-185 fully antagonized MDL induction of pCREB in both mouse OPCs (Fig. 4e-g).

#### Pharmacologic properties of GPR17 oxysterol ligands

We next sought to confirm and extend prior reports of agonism of GPR17 by certain oxysterols, most notably the brain-enriched oxysterol 24S-hydroxycholesterol (24S-OHC) and determine if other classes of lipids act as agonists of GPR17. We conducted a 1000 lipid library cAMP activity-based screen for GPR17 activity (Supplemental Tables 1 and 2). In addition to 24S-OHC, we identified two novel oxysterols as GPR17 agonists: the synthetic oxysterols NSC-12 and 20,22-dihydroxycholesterol (20,22- DHC), a naturally occurring lipid intermediate in steroid hormone biosynthesis, also referred to as OXY-16 (Fig. 5a, Supplemental Table 2), thus supporting a previous report that oxysterols are promiscuous ligands of GPR17.^13^

**Figure 5.**
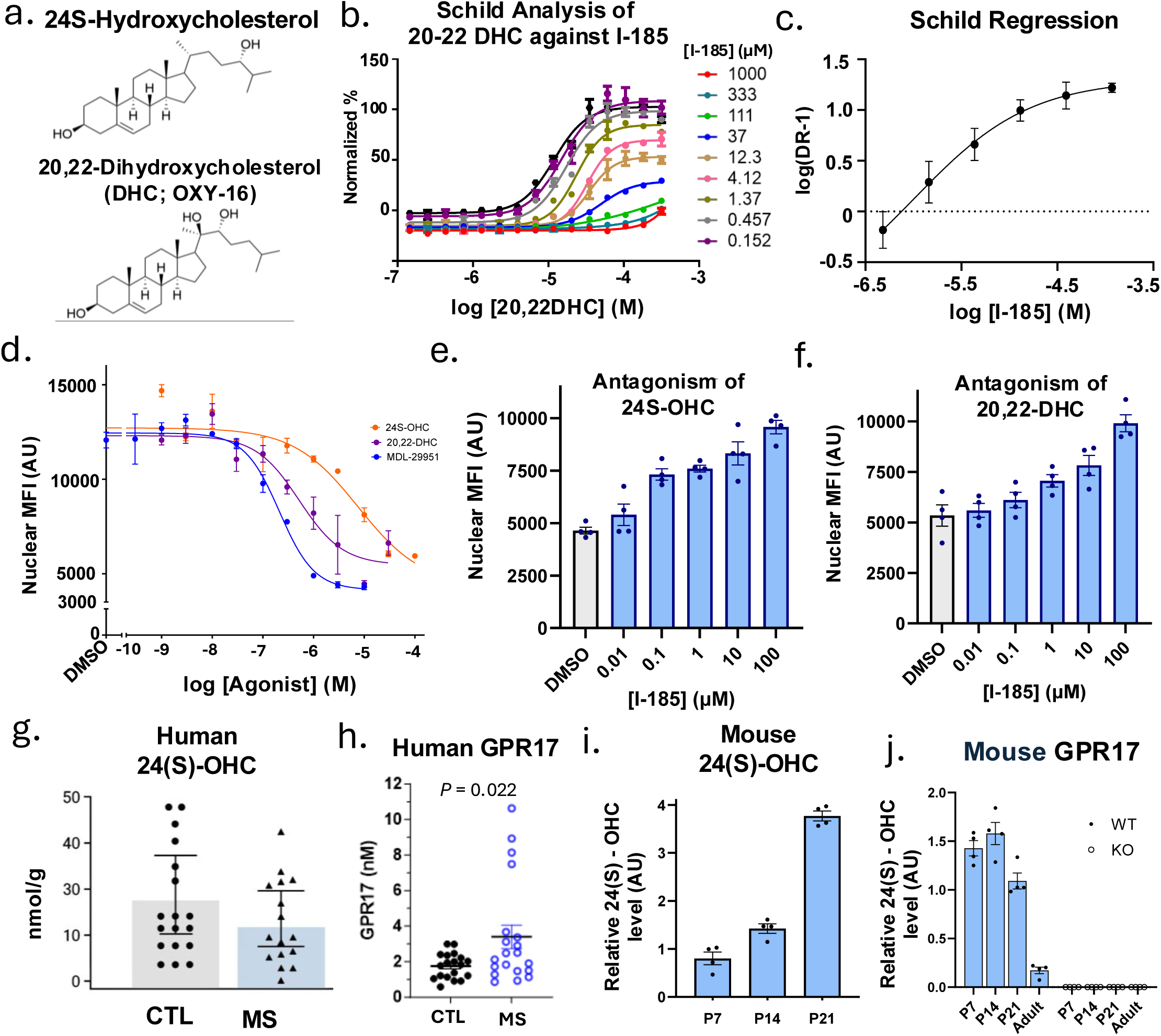
Agonism of GPR17 by select oxysterols. a) Structures of 24S-Hydroxycholesterol (24S-OHC), a previously discovered GPR17 agonist, and 20,22-Dihydroxycholesterol (20,22-DHC), a novel oxysterol agonist revealed by lipid screening. b) Schild analysis for 20,22-DHC against MDL. Normalized percentage is the change in signal from agonist relative to forskolin alone. c) Schild regression for 20,22-DHC activity. d) Dose response curves demonstrating GPR17 agonism by 24S-OHC and 20,22-DHC against MDL. e) GC-MS measurements of GPR17 over the course of myelin development in mouse (P7, P14, and P21). f) GC-MS measurements of 24S-OHC in human multiple sclerosis and control brain (cerebral white matter). g) Measurements of mouse GPR17 protein levels over the course of myelin development. h) Measurements of GPR17 protein level in control and MS brain (forebrain white matter). Plots and error bars in a to d are mean ± SEM. *N* = 4 independent experiments with 2 to 4 technical replicates. Dots in e-h represent individual brain samples. Plots and error bars are mean ± SEM.

We then characterized the pharmacologic properties of oxysterols (24S-OHC and 20,22-DHC). Owing to its poor solubility, 24S-OHC did not perform well in the saline-based HTRF assay. Therefore, we first focused on 20-22 DHC for potency, efficacy, and Schild analysis in the GPR17 astrocytoma cell line. We found that 20-22 DHC did not exhibit activity in the parent line, confirming its selectivity (data not shown). Schild analysis revealed that, in contrast to MDL, 20-22-DHC antagonism showed a nonlinear slope with E_max_ decreasing with increasing concentrations of MDL (Fig. 5c). This is a profile consistent with non-competitive binding and allostery. We next assessed activity of 20,22 DHC and 24S-OHC in human OPCs compared to MDL in the pCREB nuclear localization assay. We observed that 20,22-DHC and 24S-OHC were active agonists in human OPCs at low micromolar concentrations (Fig. 5e). Finally, we determined that I-185 could antagonize both 20,22-DHC and 24(S)-OHC (Fig. 5e,f) in a concentration-dependent manner.

#### Quantification of sterols and GPR17 protein expression in brain samples

We next sought to measure the concentrations of oxysterols in the human brain in control and multiple sclerosis patients to determine if the potencies observed in cell systems were at concentrations of potential biological relevance. We found that 24(S)-OHC was expressed at ∼20 nmol/g in control human brain and ∼17 nmol/g in chronic multiple sclerosis white matter lesions (Fig. 5g). These measurements correspond to approximate concentrations of 15 to 20 uM, in line with pharmacologically active concentrations observed in human OPC cultures. Further, we quantified GPR17 level in human brain, comparing healthy subjects and MS disease patients. In this cohort, the brain concentration of GPR17 ranged from 1-10nM and was significantly higher in MS patients (Fig. 5h).

Finally, we sought to determine the expression of both GPR17 protein and oxysterols 24S-OHC in postnatal mouse development through early adulthood; the time that correlates to the period of developmental myelination (Fig. 5i). Given that GPR17 expression is spatially and temporally specific to early OPCs, its protein abundance in whole brain is low and age dependent. Thus, we profiled its expression across development stages and we developed a parallel-reaction-monitoring (PRM)-based targeted proteomics assay using heavy isotope labeled, unique peptides of GPR17 as internal standards to achieve absolute quantification with high sensitivity and specificity. We found the GPR17 levels rose throughout brain development, peaking in the early juvenile period (P14) which corresponds to the transition from predominant OPC differentiation to myelin maturation (Fig. 5j). Conversely, 24(S)-OHC peaked later in development at P21.

#### Photolabeling and MS analysis identify competitive binding site for 24S-OHC with cholesterol

To map potential sterol binding pockets in human GPR17, we performed photoaffinity labeling experiments with a cholesterol analogue that contains a photoactivatable diazirine group and a ‘clickable’ alkyne handle. Upon ultraviolet (UV) light irradiation, activated diazirine group produces a reactive carbene that covalently modifies proximal residues and facilitates the detection of protein-lipid interaction. PhotoClick cholesterol analogs have been widely used to identify and characterize physiologically relevant sterol binding sites on membrane proteins including GPCRs in both in vitro and in cellular assays.^31,35–37^

First, we assessed photoaffinity cholesterol labeling of GPR17 using SDS-PAGE analysis. Following UV crosslinking, probe-labeled protein was conjugated to the TAMRA reporter tag by copper-catalyzed azide-alkyne cycloaddition (CuAAC or click chemistry), separated by SDS-PAGE and then visualized via fluorescent imaging (Fig. 6a). To validate the specificity of probe labeling, we examined whether the presence of excess unlabeled cholesterol could competitively block photolabeling of GPR17. Since the potency of 24S-OHC agonism for GPR17 is consistent with its physiological concentration^31^, we included 24S-OHC in our competition assay, as well as its stereoisomer (24R-OHC) to investigate the stereospecificity of lipid binding. As shown in Fig. 6, photolabeling of GPR17 is potently competed by 24S-OHC in a dose-responsive manner, while weaker competitive inhibition is observed for cholesterol (FIG. 6b). Interestingly, 24R-OHC was unable to compete with cholesterol probe labeling at a biologically relevant concentration of ∼20uM, suggesting the lipid-protein interaction is stereoselective.

**Figure 6.**
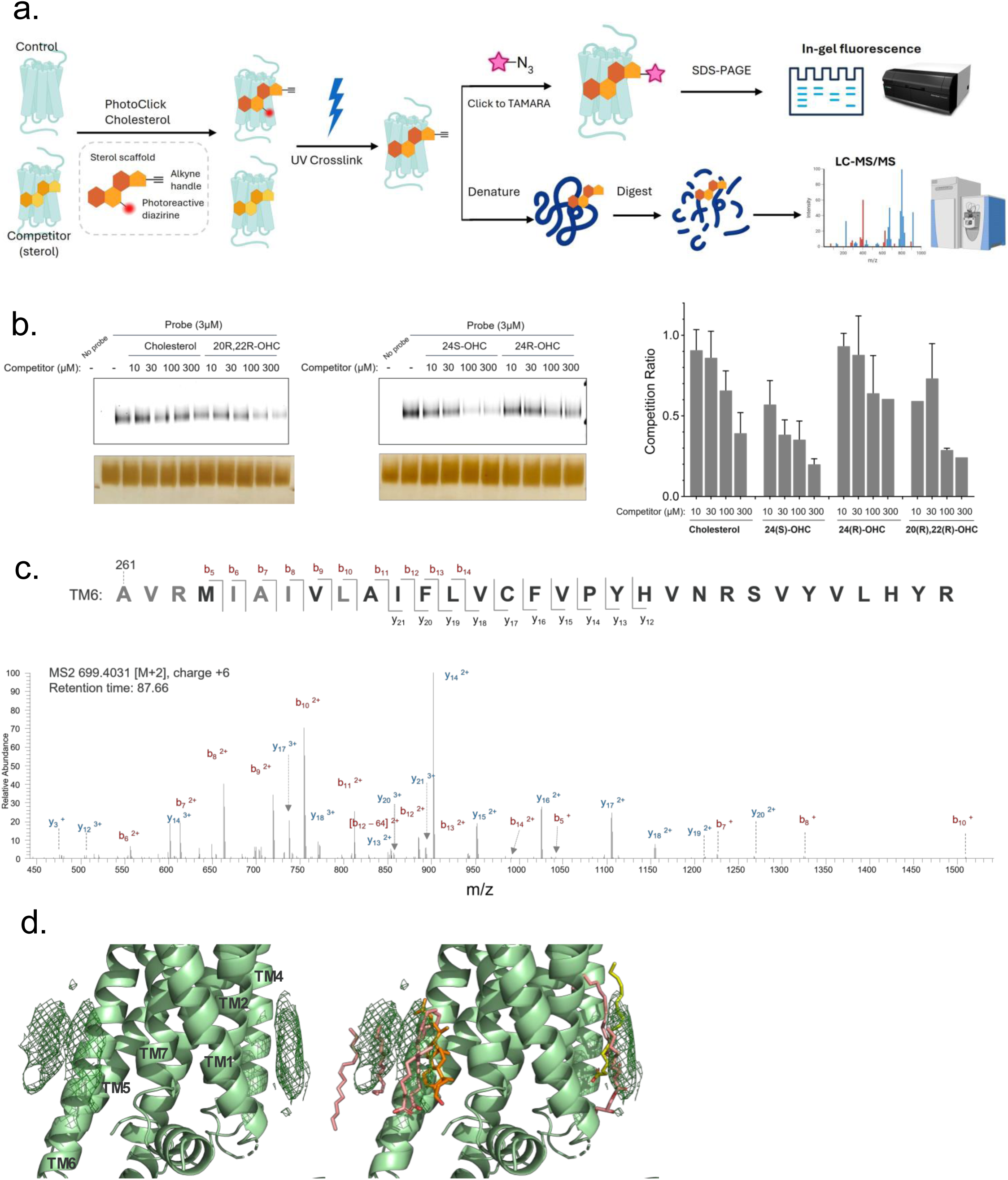
Effects and location of lipid binding to GPR17. a) Representative in-gel analysis of PhotoClick cholesterol labeling of recombinant GPR17 shown in grayscale. GPR17 was incubated with indicated lipid followed by 3µM PhotoClick cholesterol analog, crosslinked at 365 nm UV light, conjugated to TAMRA fluorophore and separated by SDS-PAGE. Photolabeled protein was imaged by Typhoon and total protein loaded in each sample was measured by silver stain. b) Competition ratio of GPR17 photolabeling by sterols at each concentration in gel-based assay. Probe labeling was quantified by normalizing fluorescent intensity (TAMARA) to protein loading (silver stain) in each lane via ImageJ. Data are mean ± SEM of *n*=4 independent replicates. c) MaxQuant database search identifying several N-terminal residues in indicated peptide are modified with a mass consistent with photolabeling (+454.344695) colored in gray. Red brackets label product ions that contain the probe adduct. Ballesteros-Weinstein numbering of residues in blue above sequence. d) Fragmentation (HCD) spectrum of the indicated photolabeled peptide mapped to TM6 (mass error: 2.5ppm). Red and blue represent b and y ion fragments that do and do not contain probe modification, respectively. e) Sterol like densities from the GPR17-MDL structure surrounding the GPR17-MDL model without (right) and overlaid with (left) lipid molecules from similar structures (orange, cholesterol from GPR161 (PDB 8KH4); yellow, oleic acid from CysLT1 (PDB 6RZ5); pink, oleic acid from CysLT2 (PDB 6RZ6).

We next set out to locate the sterol binding sites by mass spectrometry. Tryptic digests of photolabeled GPR17 were analyzed by bottom-up LC-MS/MS sequencing. A unique peptide in TM6 carrying probe adduct (+454.344) was the most prominent site identified (Fig. 6c). MaxQuant analysis reported multiple amino acids within N-terminal region of this peptide to be possible modification sites, and manual examination of its fragmentation spectrum verifies a series of *b* ions adducted with the modification site. To determine the lipid-binding specificity of identified peptide, we quantified probe-labeling efficiency in a competition assay. We observed a stronger competitive inhibition of photoprobe labeling by 24S-OHC, compared to cholesterol, which agrees with our in-gel fluorescence analysis. Taken together, our photolabeling data locates a lipid binding pocket in GPR17 with specificity towards 24S-OHC.

#### Sterol-like densities in GPR17 Structure

In solving the structure of MDL-bound GPR17 we identified several potential sterol-like densities (SLD) surrounding the transmembrane domain (TMD) (Fig. 6d, SLD1-4). When we compared the agonist-bound map to the apo map (PDB 7Y89), some of these densities were also present. This suggested that these were not artifacts from data processing, but potential ligand-binding sites of interest, especially as it has been hypothesized that >5 different oxysterols can activate GPR17.^13^ One primary peptide from TM6 was shown to be modified by the probe by mass spectrometry analysis, which identified a potential allosteric binding site for cholesterol in comparison to the previously predicted orthosteric binding mode. There are at least three SLDs that flank TM6, making this observation feasible, however they are at a distance from the TMD >4 Å, limiting definition of any specific direct interactions like those shown for GPR161, which are as close as 3 Å to V274^6.39^ and 3.4 Å to W327^7.56^. Notably, GPR17 SLD1 closely aligns with the cholesterol in GPR161 and with oleic acids in the CysLT1 and CysLT2 structures. The largest unassigned density in the MDL-GPR17 structure, SLD4, aligns with a known GPCR cholesterol binding site reported in multiple structures along the exterior of the TM2-TM3-TM4 interface. The density is also present in the antagonist structure, though less significant. There are also oleic acid molecules in these SLD locations in both CysLT2 and CysLT1 antagonist-bound structures, which were solved by x-ray diffraction in the lipidic cubic phase.

## Discussion

Our studies identify the binding sites and modes of 2 known modulators of GPR17 to enable drug discovery and describe mechanisms of GPR17 activation. The ligand-bound cryo-EM structures of GPR17 in either the active or inactive state show two critical things: 1) the ligands occupy the side of the orthosteric site near the TM4-TM5 proposed sterol access site identified in related receptors like EBI2, and 2) the ECL2 is effectively fixed in its “self-activating” position in active, inactive, and apo states. Together this suggests that druggability of the receptor is limited by an atypically small pocket with restrictive access. This structural data, combined with the binding and pharmacology data herein, suggest that the select oxysterols that modulate the receptor do so stereoselectivity from an allosteric site.

In this work we defined the binding sites of two GPR17 modulators: MDL agonist and I-185 antagonist. Both molecules bind in the orthosteric site, which is significantly reduced in size due to the consistent occupation of this pocket by the ECL2 in all structures solved thus far. The compact orthosteric site exhibits a strong positive charge and provides a vector out of the entry site between TMs 4 and 5 that leads into a hydrophobic channel on the membrane-embedded surface of GPR17. Despite ligand similarities in indole substituents, their placement and interactions have subtle differences that enable the activation state of GPR17, specifically via shifts in bulky hydrophobic residues (Y140^3.33^, Y144^3.37^, Y148^3.41^) on TM3 that cascade to the transmission switch (I147^3.40^). Observation that the ECL2 of GPR17 remains locked in place allows reinterpretation of previous SAR findings in the context of experimental structures. Our structures correlate with the steep SAR findings for both agonists and antagonists that the compound must have a central strong electronegative feature to interact with positively charged orthosteric site residues (ex. R283^6.55^, R214^5.32^, H220^5.38^, H280^6.52^), and variable substitution is only tolerated in particular locations around the ring system, specifically locations that lead into the hydrophobic groove (ex. Y140^3.33^, Y144^3.37^, L194^4.60^, L225^5.40^). Penetration of this shallow hydrophobic groove is necessary for improved potency. The shared constraints on variable substitutions allowing penetration of this shallow hydrophobic groove make differentiating agonists and antagonists difficult and therefore may obfuscate rational design of inhibitors versus activators.

The ECL2 of GPR17 is stable in its position which is thought to lead to self-activation, yet there is structural and pharmacological data confirming subtle GPR17 modulation by synthetic ligands. The interactions between Y213^ECL2^ and R308^7.36^ or H119^2.64^, R214^ECL2^ and N307^7.35^, and the C132^3.25^-C209^ECL2^ disulfide bond are present in apo and both ligand-bound structures. Structural studies of orphan, self-activated GPCRs have shown that they contain ECL2s that block access to the orthosteric site (BILF2, GPR61) and/or occupy the orthosteric site providing the interactions for activation (GPR52, GPR21, GPR161); however, there are very few structures available to understand how ligands might manipulate their activity (Extended Data Fig. 6)

Our structural analysis and competition experiments suggest that an activator like 24S-OHC is unlikely to fit in the GPR17 orthosteric site while simultaneously occupied by ECL2, despite the presence of a similar sterol agonist (7a,25-dihydroxycholesterol, DHC) in the orthosteric site of EBI2. Indeed, the placement of the GPR17 agonist MDL overlays well with the placement of the heterocyclic core of DHC in EBI2, but the aryl tail of DHC occupies the same space as the GPR17 ECL2. Moreover, our pharmacological inhibition data shows that select activating sterols do not compete directly with MDL. This is in line with a previous observation that potentiation of MDL by oxysterol could be fully inhibited by a GPR17 antagonist. Our pharmacological and binding data confirm and extend these observations demonstrating a Schild regression profile consistent with noncompetitive binding of the GPR17 antagonist I-185 and the oxysterol 20,22-DHC. This is in contrast to MDL, which exhibits properties indicative of direct competition with I-185. We further suggest that the proposed native ligand 24S-OHC, which we show does not compete with agonist small molecule MDL, modulates the receptor from an allosteric site in proximity to TM6. Interestingly, this activation of GPR17 by sterols appears stereo-selective, as we show that 24S-OHC but not 24R-OHC can outcompete a labeled cholesterol molecule and we identify additional activating sterols from a cAMP -based screen for GPR17 activity. The proposed placement near TM6 is similar to that of the cholesterol shown to modulate GPR161 activity, which has direct interactions with TM6 within 3Å and is thought to modulate the stability of the GPR161 - G_αs_ protein heterotrimer complex. Additional studies will be required to narrow the specific binding site on GPR17 and identify the stabilizing interactions of these specific sterols, however the presence of SLDs in the cryo-EM maps of the agonist-bound structure in the area flanking TM6 and the identification of a peptide from TM6 in the MS-labeling experiment suggest a potential specific allosteric site. Further elucidation of the allosteric site for oxysterol binding and identification of antagonists that directly compete with or modulate the allosteric site offer a compelling new avenue for GPR17 antagonism. This approach would also directly leverage the emerging understanding of oxysterols as potential natural ligands for GPR17.^13^

The findings that 24S-OHC is an agonist for GPR17 in human OPCs and expressed at around 18 nmol per gram tissue (approximately 17 µM) in human controls and multiple sclerosis brain provides further support, and the first human evidence, that 24S-OHC might be a biologically relevant ligand for GPR17. The correlation of 24S-OHC and GPR17 during the developmental period of myelination further suggests a role for 24S-OHC mediated GPR17 regulation in oligodendrocyte maturation. The concentrations of both GPR17 and 24S-OHC rise and peak into the early juvenile period.^13^ In adulthood, GPR17 expression drops substantially while 24S OHC remains elevated. This shift in the ratio of 24S-OHC to GPR17 concentration may regulate the limited OPC differentiation and myelin remodeling that occurs in adulthood.^38–40^ However, as previously discussed^13^, hypothesized roles for 24S-OHC in regulation of myelination and remyelination must be tempered and contextualized until the free fraction of 24S-OHC is determined. Ideally, these uncertainties will be addressed by further advances enabling quantification of unbound 24-OHC combined with experiments that modulate production of 24S-OHC in models of myelin development and remyelination.

In sum, our findings advance our understanding of the structural pharmacology of GPR17 and ligand receptor relationships with implications for therapeutic discovery. By resolving the structural basis of orthosteric antagonist versus agonist ligands, we reveal that GPR17 has a small binding pocket constrained by a fixed self-activating ECL2. These properties provide a potential explanation for the limitations in GPR17 druggability evidenced by the relatively narrow chemical space that typifies the sulfonamide scaffolds of many published GPR17 antagonists. The structural details of GPR17 antagonism may guide the design of improved and differentiated GPR17 antagonists targeting this challenging binding pocket. Alternatively, our characterization of allosteric regulation of GPR17 by oxysterols presents a potential framework for GPR17 antagonism with allosteric modulators.

## Methods

### Structure

#### Construct design and protein expression for cryo-EM study

The expression construct for human GPR17 in the active conformation (KC165) was modified from wild type with a haemagglutinin signal peptide, twin-strep tag, N-term addition of β2-Adrenergic receptor, and HRV-3C cleavage site, before the full length 1-367 GPR17 with C-terminal TEV cleavage site, and 10×histidine and Avitags.

The expression constructs for human GPR17 in the inactive conformation (KC185, 221) were designed with an HA signal peptide, FLAG and twin-strep tags, N-term addition of β2AR, HRV3C cleavage site, truncated GPR17 (A48-K343) followed by TEV and 10x His-tag, with bRIL insertion at G250 and L257, with E and V residues added as linkers that align to the linkage in the CysLT2-BRIL fusion (PDB 6RZ6^25^) (KC185); Q249 and L257, with modified linkers from the A_2A_ adenosine receptor (ARRQL between TM5 and N-terminus of BRIL, and ERARSTLV between C-terminus of BRIL and TM6)^22,24,41^ (KC221). Both had D105^2.50^N and F158^3.51^Y mutations.

Additional constructs to compare bRIL insertion and mutation effects KC225 were designed as the wild-type active construct, with the bRIL insertion at Q249 and L257, with modified linkers from the A_2A_ adenosine receptor. Edits were made by Q5 site directed mutagenesis (NEB).

GPR17 constructs were synthesized (Azenta) and subcloned with restriction sites (EcoRI/BamHI) or HiFi assembly (New England Biolabs) into a BacMam dual vector with the polyhedrin promoter followed by the CMV enhancer and promoter, the T7 promoter, the gene of interest and C-terminal WPRE element 5’ to 3’, and VSV-G under the p10 promoter was included in the reverse direction 3’ to 5’.

BacMam constructs were transformed into MaxEfficiency DH10Bac (Gibco) competent cells and isolated colonies post blue-white selection were cultured for isolation of bacmid DNA, which was transfected with Cellfectin II Reagent (Gibco) into Sf9 cells for virus generation. Virus was amplified for 2 passages, then used to infect HEK293SGnTi^-^ cells (ATCC CRL-3022) in suspension at 10% (v/v).

Cells were grown at 30°C and enhanced with 10 mM sodium butyrate for 64 h, then harvested by centrifugation and flash frozen at −80°C prior to membrane preparation. Expression was confirmed by western blot.

#### GPR17 Membrane preparation

HEK293SGnTi^-^ cells were lysed in low salt buffer containing 10 mM HEPES pH 7.5, 10 mM MgCl_2_, 20 mM KCl supplemented with benzonase and Pierce protease inhibitor tablet, dounced, and ultracentrifuged for 30 min at 45,000 rpm at 4°C. Pellets were then dounced in high salt buffer containing 10 mM HEPES pH 7.5, 10 mM MgCl_2_, 1M NaCl, 20 mM KCl supplemented with high salt benzonase and Pierce protease inhibitor tablet and ultracentrifuged. Membranes were dounced in low salt buffer containing 20% Glycerol, and flash frozen at −80°C.

#### Purification of SapA

Human Saposin A (SapA) expression was performed as previously described.^42^ Briefly, the coding sequence for SapA was synthetized (Azenta) and subcloned into a pET28a vector backbone with N-terminal 8x histidine tag and TEV protease cleavage site. The protein was expressed using *E. coli* Rosetta gami-2 cells (DE3)(Novagen). For expression, pre-cultures were set up by inoculating Terrific broth (TB) medium supplemented with 12 ug/mL Tetracycline and 50 ug/mL Kanamycin with single colonies and incubated over night at 37°C with shaking. The main culture was inoculated with antibiotics at 10 mL/L in TB. At OD_600_ ≈ 0.6-0.8, protein expression was induced with 0.7 mM IPTG, and expression continued at 25°C for 16-20 h. Cells were collected by centrifugation and the cell pellet was stored at −80°C.

For protein purification, the cell pellet was thawed and resuspended in 20 mM HEPES pH 7.5, 150 mM NaCl, 20 mM Imidazole pH 7.5 supplemented with benzonase, and lysed by high pressure homogenization (Microfluidics). Lysates were subjected to centrifugation at 40,000 xg, followed by heating of the supernatant for 10 min at 85°C and 300 rpm, followed by another centrifugation step. The supernatant was loaded onto a pre-equilibrated HisTrap FF crude column (Cytiva) at 2 mL/min and eluted by stepwise gradient with buffers containing 20 mM HEPES pH 7.5, 150 mM NaCl and various concentrations 20-500 mM Imidazole pH 7.5 with SapA eluting at 400 mM Imidazole. The monomeric peak fractions were pooled, supplemented with AcTEV protease (Invitrogen) and 1 mM DTT, and dialyzed overnight against 20 mM HEPES pH 7.5, 150 mM NaCl. Cleavage was confirmed by SDS gel. TEV protease and the cleaved histidine tag were removed by Reverse-IMAC, and the flow-through was subjected to size exclusion chromatography using HiLoad Superdex 75 16/600 GL (Cytiva) using the dialysis buffer as running buffer. Peak fractions were pooled, flash-frozen in liquid nitrogen and stored at −80°C. Typically, yields of 10-15 mg/L were obtained.

#### Purification of GPR17 agonist structure construct

Membranes containing WT GPR17 (KC165) were thawed and resuspended in buffer containing 50 mM HEPES pH 8, 500 mM NaCl, 5 mM MgCl_2_, 2 mg/mL iodoacetamide and 5 µM MDL and rocked for 1 h before the addition of 2x solubilization buffer containing 2% lauryl maltose neopentyl glycol (LMNG, Anatrace) and 0.2% cholesterol hemisuccinate (CHS, Anatrace), 50 mM HEPES pH 8, 1M NaCl, 5 µM MDL, and spun at 4 °C for 3 h. The supernatant was clarified by centrifugation and batch-bound to StrepTactinXT resin (IBA Lifesciences) overnight. The resin was washed with 10 column volumes of wash I buffer (50 mM HEPES pH 8, 1M NaCl, 5 µM MDL, 5 mM MgCl_2_, 5 mM ATP, 0.1% LMNG, 0.01% CHS), then 10 column volumes of wash II buffer (wash I buffer without ATP added) and eluted with 50 mM HEPES pH 8, 1M NaCl, 5 µM MDL, 5 mM MgCl_2_, 0.05% LMNG, 0.005% CHS, 50 mM biotin. Elution was batch-bound to washed Talon resin (Takara) for 1 h in the presence of 10 mM imidazole, washed with 50 mM HEPES pH 8, 800 mM NaCl, 5 µM MDL, 5 mM MgCl_2_, 0.025% LMNG, 0.0025% CHS, 20 mM imidazole, and eluted with 50 mM HEPES pH 8, 800 mM NaCl, 5 µM MDL, 5 mM MgCl_2_, 0.0125% LMNG, 0.00125% CHS, 300 mM imidazole. The sample was concentrated in a 50 kDa MWCO Amicon filter before size exclusion chromatography on Superdex 200 Increase 10/300 GL column (Cytiva) with 50 mM HEPES pH 8, 500 mM NaCl, 5 µM MDL, 5 mM MgCl_2_, 0.0125% LMNG, 0.00125% CHS running buffer. The monomeric fraction was subjected to HRV3C cleavage to remove affinity tags overnight, then reverse IMAC was performed the following day before a second size exclusion step completed in 50 mM HEPES pH 8, 500 mM NaCl, 5 µM MDL, 5 mM MgCl_2_, 0.0125% LMNG, 0.00125% CHS running buffer. Fractions containing cleaved, monomeric GPR17 were used for further analysis and agonist structure determination after combination with G protein heterotrimer. For the complex, the protein was combined with equimolar purified Gα_i_ protein heterotrimer in the presence of 25 mU/mL of apyrase (New England Biolabs) and 2x Molar scFv16 (Thermo Fisher) at 4 °C for 1 h, then purified by size exclusion on a Superdex 200 Increase 10/300 GL column (Cytiva) with 20 mM HEPES 8, 250 mM NaCl, 0.00075% LMNG/CHS, 0.00025% GDN, 5 µM MDL (MedChem Express) running buffer. A single, 0.5 mL aliquot containing the complex was concentrated for grid preparation.

#### Purification of GPR17 Antagonist structure constructs with SapA

Membranes of modified GPR17/KC221 in inactive conformation were thawed and resuspended in buffer containing 50 mM HEPES pH 7.5, 500 mM NaCl, 5 mM MgCl_2_, 2 mg/mL iodoacetamide and stirred for 1 h at 4°C. Solubilization was performed by addition of 2x solubilization buffer containing 1.5% lauryl maltose neopentyl glycol (LMNG, Anatrace) and 0.15% cholesterol hemisuccinate (CHS, Anatrace), 100 mM HEPES pH 7.5, 1 M NaCl, and incubated at 4 °C for 3 h. The supernatant was clarified by ultracentrifugation for 1h at 45,000 rpm at 4°C (Ti-45, Beckman Coulter) and used for nanodisc reconstitution.^43^ Briefly, the supernatant was mixed with purified SapA in a concentration of 3 mg_SapA_/L_cells_ and incubated for 30 min at 4°C rotating. To remove excess detergent, Bio-Beads SM-2 Resin (Bio-Rad) was added to the solution in a concentration of 2.5 g_Bio-Beads_/L_cells_, and batch-bound to StrepTactinXT resin (IBA Lifesciences) overnight. The resin was washed with 10 column volumes of detergent-free wash buffer containing 50 mM HEPES pH 7.5, 800 mM NaCl, 5 mM MgCl_2_ and two times incubated for 30 min with 50 mM HEPES pH 7.5, 500 mM NaCl, 5 mM MgCl_2_, 50 mM biotin and 0.05 mg/mL SapA for elution. Elution fractions were batch-bound to washed Talon resin (Takara) for 1 h, washed with wash buffer, and eluted with 50 mM HEPES pH 7.5, 500 mM NaCl, 5 mM MgCl_2_, 300 mM imidazole pH 7.5, and 0.0625 mg/mL SapA. The sample was concentrated in a 50 kDa Amicon MWCO filter before size exclusion chromatography on Superdex 200 Increase 10/300 GL column (Cytiva) with 50 mM HEPES pH 7.5, 500 mM NaCl, 5 mM MgCl_2,_ 1 mM TCEP as running buffer. For the antagonist complex, the purified GPR17-SapA complex was combined with 1.3 molar excess of purified anti-bRIL Fab, 1.5 molar excess of purified anti-Fab Nb, and 100 µM antagonist I-185 (Enamine), and incubated over night at 4°C rocking. The complex was purified by size exclusion on a Superdex 200 Increase 10/300 GL column (Cytiva) with 50 mM HEPES 8, 250 mM NaCl, 5 mM MgCl_2_, 10 µM I-185 as running buffer. The peak fractions were collected and concentrated to 2-3 mg/mL. The final protein sample was supplemented with I-185 to a final concentration of 100 µM and subjected to structure determination.

#### Purification of G protein heterotrimer

G protein heterotrimer was expressed and purified separately from GPR17, from co-expression of human Gα_i1_, Gβ_1_ and Gγ_2_. Open reading frames were synthesized and subcloned in a modified pFastBac1 vector (Thermo Fisher) for expression in Sf9 cells. The Gα_i_ construct was the dominant-negative Gα_i1_ with G203A and A326S stabilizing mutations with reduced nucleotide binding affinity.^44,45^ Gβ_1_ contains an N-terminal leader of MERK-6×histidine tag-GSSG- before amino acids S2-N340 (UniProt P62873) and Gγ_2_ corresponds to M1-C71 (UniProt P59768). pFastBac constructs were transformed into MaxEfficiency DH10Bac competent cells to generate virus according to the Bac-to-Bac system protocol (Thermo Fisher). Sf9 cells at a cell density of 3.5×10^6^ cells/mL were infected with P2 virus at 1%/0.5%/0.5% (v/v) Gα_i1_/Gβ_1_/Gγ_2_, harvested after 48 h and frozen at −80 °C. Cells were lysed in 10 mM HEPES pH 7.5, 10 mM MgCl_2_, 5 mM TCEP, 50 µM GDP, Pierce protease inhibitor cocktail (1 piece/50ml) by dounce homogenization, then centrifuged. Membrane fractions were dounced in solubilization buffer: 50 mM HEPES pH 7.5, 150 mM NaCl, 5 mM TCEP, 5 µM GDP, 1% LMNG, 0.2% CHS, 10 mM imidazole, and stirred for 3 h before centrifugation. The solubilized fraction was passed over a 5 mL HiTrap Talon Crude column (Cytiva) pre-equilibrated in buffer A: 50 mM HEPES pH 7.5, 150 mM NaCl, 5 mM TCEP, 5 µM GDP, 0.1% LMNG, 0.02% CHS, 20 mM imidazole, washed with buffer B: 50 mM HEPES pH 7.5, 150 mM NaCl, 5 mM TCEP, 5 µM GDP, 0.01% LMNG, 0.002% CHS, 50 mM imidazole, and eluted with buffer C: 50 mM HEPES pH 7.5, 150 mM NaCl, 5 mM TCEP, 5 µM GDP, 0.005% LMNG, 0.001% CHS, 250 mM imidazole. The eluted sample was concentrated and loaded on a Superdex200 10/300 pre-equilibrated with SEC buffer: 50 mM HEPES pH 7.5, 150 mM NaCl, 10% glycerol, 5 mM TCEP, 5 mM GDP, 0.003% LMNG, 0.0006% CHS. Fractions containing the heterotrimer, identified by SDS PAGE, were pooled and concentrated to < 2 mg/mL. (Extended Data Fig. 1)

#### Purification of anti-bRIL Fab and anti-Fab Nb

Synthetic anti-BRIL Fab was expressed by transient transfection in HEK CHO cells using vectors for heavy and light chains with the chemotype as described.^46^ A DNA fragment encoding the anti-Fab nanobody (Nb) was synthesized (Azenta) and cloned in pET28a vector with an N-terminal pelB signal sequence followed by a 6×His tag and TEV protease site and was purified as described.^47^ ^48^

#### Cryo-EM sample preparation

For cryo-EM, 3 µL aliquots of GPR17 complex were applied to grids that had been glow discharged for 120 s at 20 mA. For the MDL-bound structure, SEC-purified complex at ∼1.3 mg/mL and R 0.6/1, Au, 300 mesh Quantifoil grids were used. For the I-185-bound structure, 0.025% A8-35 was added to SEC-purified complex at ∼1.4 mg/mL then R1.2/1.3, Au, 300 mesh UltrAuFoil grids were used. Using a Vitrobot system (Thermo Fisher Scientific), samples were applied, then grids were immediately blotted for 5s at blot force 7 at 4 °C and 100% humidity before plunge freezing and subsequent vitrification in 100% liquid ethane. Before screening and data collection, grids were clipped into autogrid assemblies.

#### Cryo-EM data collection and processing

During grid optimization, grids were screened in-house using SerialEM on a Glacios electron microscope (Thermo Fisher Scientific) system with a K3 direct electron detector (Gatan). Grids resulting in promising 2D classes were taken to the MIT Characterization.nano facility for high-resolution data collection. The final data set was collected using the EPU software (Thermo Fisher Scientific) on a Titan Krios G3i (Thermo Fisher Scientific) with a K3 camera (Gatan). For the MDL-bound structure, 10,712 movies were collected in a defocus range of −0.25 to −2.0 at ×130,000 in superresolution mode with a pixel size 0.663 Å/pixel and a total dose of 65.712 e−/Å^2^ per micrograph. For the I-185-bound structure, 15,507 movies were collected in a defocus range of −0.25 to −2.5 at ×130,000 in superresolution mode with a pixel size 0.654 Å/pixel and a total dose of 62.484 e−/Å^2^ per micrograph. Initial motion correction and CTF estimation were performed in cryosparc live.^49,50^ ^51^. Template Picking, Remove Duplicates, Ab-Initio, Heterogeneous Refinement, Homogeneous Refinement, Global CTF Refinement, and Non-Uniform Refinement) in cryosparc (Extended Data Fig 2). Density modification of the final map for the MDL-bound structure only was performed in Phenix Resolve.^52^

#### Model building, refinement, and comparison

To generate the model of GPR17 in complex with MDL, the publicly available AlphaFold (https://doi.org/10.1038/s41586-021-03819-2) model (AF-Q13304-F1-v4) and the heterotrimeric complex model from Cannabinoid Receptor 1-G Protein Complex (PDB 6N4B)^53^ were fit into the experimental map using USCF Chimera 1.18,^54^ manually rebuilt (including ligand building and placement) in Coot ^55^ and refined (real-space) in Phenix.^52^ The model of GPR17 in complex with I-185 was created from our in-house structure with MDL and rebuilt and refined as above. Figures were made in PyMOL (The PyMOL Molecular Graphics System, Version 3.0, Schrödinger, LLC.). Any compared model files were aligned in PyMOL or Coot based only on the 7TM helix domain. Maps were compared over the 7TM domain in Chimera. UCSF Chimera was developed by the Resource for Biocomputing, Visualization, and Informatics at the University of California, San Francisco, with support from NIH P41-GM103311.

### In vitro pharmacological assays

#### Cell line generation and cAMP detection in 1321N1 cell line

Human astrocytoma cells (1321N1) stably expressing GPR17 (1321N1-GPR17) were used to measure intracellular cAMP levels after compound treatment using Revvity’s cAMP displacement HTRF (homogeneous time resolved fluorescence) kit. 1321N1-GPR17 cells were cultured in DMEM, 10% FBS, 1x P/S, under 5 µg/mL puromycin selection at 37 °C, 5% CO_2_. Cells were lifted from TC flask by washing with 1x PBS followed by trypsinization for 5 min. Cells were pelleted, resuspended in Opti-MEM and counted. Cells were diluted to 0.7×10^6^ cells/mL in Opti-MEM containing 0.5 mM IBMX, 10 µL was added to a white 384 well TC treated plate. The plates were sealed and spun for 1min at 1000 rpm and incubated at room temp for 15min. Cells were stimulated with serially diluted MDL with 2.5 µM Forskolin (final concentration) in Opti-MEM. For the Schild regression analysis the antagonist I-185 was titrated in the 384well TC treated plates via Echo prior to adding cells and the stimulated with MDL and Forskolin. The plates were sealed and spun down for 1 min at 1000 rpm and incubated at ambient temperature for 15 min. 10 µL of 1x cAMP-d2 in lysis buffer was added and spun down, followed by addition of 10 µL of 1x Anti-cAMP Eu cryptate antibody in lysis buffer (for 0.25x cAMP-d2 and Anti-cAMP Eu cryptate final concentration). Plates were sealed, spun 1 min at 1000 rpm, incubated at ambient temperature for 1 hr, then read on Perkin Elmer Envision using the HTRF ratio based raw data at 665 nm/ 615 nm wavelength.

For data analysis the readout value (10^4^ x 665 nm/ 615 nm) for each compound was normalized with the value of the 0% activity control and 100% activity control the same plate.

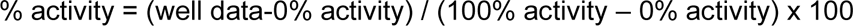

Normalized values were plotted as activity vs. dose and fit to the following 4-parameter logistic model in Graphpad Prism. The Schild regression analysis was done by plotting the log (dose ratio − 1) vs. log[antagonist] of different concentrations of I-185 in the presence of increasing concentration of MDL or 20,22-DHC. ^56^ I-185 curves that showed significant separation (students one-way T-test) from the max and min concentrations of ligand were selected for Schild analysis and regression plotting.

#### Human oligodendrocyte cultures

For culture of human IPSC-derived oligodendrocytes, transcription factor-driven ioOligodendrocytes were obtained from bit.bio (bit.bio1028) according to the manufacturer’s instructions. Briefly, cells were thawed and seeded on a 12 well or 96 well plate coated with laminin and poly-d-lysine at densities of 100,000 and 30,000 cells per well respectively. According to the bitbio user manual, cells were cultured for 24 h in M1 media containing 5 μm of the ROCK inhibitor Y-27632 (Abcam #ab144494) to stabilize the reconstituted culture and then switched to M2 media for oligodendrocyte differentiation. Doxycycline (1 μg/mL) for Tet induction of pro-differentiation transcription factors was added at the time of seeding. For cAMP and pCREB assays, Cell were cultured for 5 days in M2 media, which was determined to be a time point of GPR17 enrichment. For cAMP assays, cells were plated on polystyrene microplates with black walls (Corning #3842), for pCREB assays cells were plated on PhenoPlates for high content imaging (Revvity 6005050), and for qPCR cells were plated on clear polystyrene 12 well plates (Corning 3336).

#### GPR17 KO mouse generation and animals for primary mouse cultures

The Biogen Institutional Animal Care and Use Committee (IACUC) approved all mouse care, breeding, husbandry, and manipulations described herein under Biogen IACUC protocol 871. Animals were housed in an American Association for Laboratory Animal Science-certified facility and monitored daily by animal staff.

The GPR17 knockout mice were generated via CRISPR-Cas9 mediated excision of exon 2 and bred onto the C57BL/6J background for 6 generations. Absence of GPR17 expression was confirmed by immunohistochemistry and qPCR for GPR17 (see below for methods). Primary cultures were obtained from neonates (P6-P8) of either GPR17KO or C57BL/6J wild type mice (Jackson Laboratory Cat# 000664).

#### Primary mouse oligodendrocyte cultures

Primary mouse oligodendrocytes were obtained for P6-P8 mouse using the Miltenyi MACs kit and protocol for magnetic isolation and cultivation of mouse oligodendrocytes according to the manufacturer’s instructions (Miltenyi Biotec 130-092-628 and 130-094-543). Briefly, after isolation, cells were plated at 20,000 cells/well on 96 well PhenoPlates for pCREB high content imaging assays, expanded for two days in the presence of PDGF, and then differentiated to GPR17 expressing pre-oligodendrocytes in the presence of 600 ng/mL of T3 (MilliporeSigma #T2877).

#### cAMP kinetic functional assays in human OPCs

Functional assays for cAMP measurement were made using the BACMAM viral delivery of a cAMP sensor using the Green Gi cADDIs CAMP assay kit with CMV promoter (Montana Molecular #X0200G) according to the manufacturers protocol. Briefly, on day four of differentiation in culture, cells were transduced by treating cells incubated in 100 µL media per well with 50 µL of transduction reaction mix (containing 20 µl cADDis cAMP sensor, 0.6 µL of 500 mM sodium butyrate, and 29.4 µL media) for 30 min at ambient temperature. After 30 minutes of treatment, a full media change was made, cells were returned to the incubator, and cAMP assays were conducted 24 hours later.

Assays were conducted using real time fluorescence detection on a Perkin Elmer Envision 2014 multimodal plate reader. Individual reads were spaced by one minute. For assays of GPR17 agonism by MDL, 6 baseline reads were conducted, Forskolin was added at concentrations ranging from 0.1 to 3.0 µM to induce cAMP production downstream of Gi signaling, then ten reads were taken over ten minutes which corresponded to the time to plateau in the FSK signal. MDL was then added at concentrations ranging from 0.1 to 3.0 µM, resulting in a reversal of the forskolin signal.

The MDL response was quantified as a percent effect ratio by obtaining the ratio of the peak MDL response versus the peak FSK response. For antagonism, titrations of I-185 were preloaded before the assay. Antagonism was quantified by the percent difference in percent effect of the MDL response with vehicle versus the difference in MDL response with each concentration of I-185. The percent effect of MDL agonism and inhibition by I-185 were plotted in GraphPad Prism version 9 or 10. The half maximal effective concentration (EC_50_) of MDL agonism and the 50% inhibition of the maximal effective concentration of MDL agonism by I-185 (IC_50_) were determined using a four-parameter logistic dose response equation.

#### pCREB Nuclear Localization Assays

For mouse OPC O4+ cells, the PhenoPlate-96 were pre-coated with poly-D-lysine, and cells seeded at a density of 20,000 cells/well in Differentiation Media plus CNTF (Miltenyi Biotec, 130-096-336, 10 ng/mL). Differentiation Media consisted of DMEM (high glucose) supplemented with 1X N-2 supplement, 1X B-27 supplement, 1 X Pen-Strep, 2 mM GlutaMAX, 1 mM sodium pyruvate, 5 mg/mL insulin, 5 mg/mL N-acetyl-L-cysteine, 1X Trace Element B, 10 ng/mL d-biotin). No forskolin or db-cAMP was added to the media. The next day, media was completely exchanged and the assay was performed on day 2 post seeding. For human IPSC-derived oligodendrocytes, ioOligodendrocytes were prepared and cultured as described in the earlier human oligodendrocyte cultures subsection.

On the day of the assay, cells were pretreated for 20 – 30 minutes with compound at 37°C, 5% CO_2_. For mOPC O4+ cells, IBMX (100 – 500 mM) was included at all steps. This was followed by a 30-minute stimulation at 37°C, 5% CO_2_ (forskolin and MDL final concentrations indicated in Fig. 4).

Following stimulation, cells were fixed at 4% paraformaldehyde for 15 minutes and permeabilized/blocked in PBS containing 5% normal goat serum and 0.01 %Triton X-100 for a minumum of 60 minutes at room temperature. Antibody to Phospho-CREB (Ser133) (87G3) (Cell Signaling Technologies, 9198) was diluted 1:800 in permeabilization/block buffer (5% NGS, 0.01% Triton X-100 in PBS) and cells were stained with primary antibody overnight at +4°C. The following day, plates were washed and secondary antibodies were diluted 1:500 in PBS containing 1% NGS and 0.01% Triton X-100 and used at room temperature for 60 minutes with Alexa Fluor 647 anti-rabbit (Thermo Fisher, A-21245). Hoechst 33342 at 1:2000 was included in the secondary stain to identify nuclei. Plates were washed and data were acquired with the Operetta CLS (Revvity) at x20 water objective. Image quantification was performed using Columbus software (Revvity) which included nuclei detection, live cell identification (size and Hoechst intensity), and measurement of nuclear pCREB signal. An algorithm was developed using Columbus software to quantify 1) the media fluorescence intensity on a cell-by-cell basis and 2) the percent of pCREB+ cells on a threshold basis against vehicle treated cells. Both methods were used to determine antagonist and agonist potency.

### TaqMan Real-Time PCR for oligodendrocyte lineage marker gene detection in bit.bio

#### oligodendrocytes

RNA was harvested from bit.bio oligodendrocytes plated at 100,000 cells per well density at 1, 3 5, and 7 days of differentiation using the Zymo quick RNA mini-prep kit according to the manufacturer’s instructions (Zymo R1055).Transcript levels of the following oligodendrocyte differentiation marker genes were determined using the TaqMan qPCR platform: GPR17 (ThermoFisher probe Mm01159800_s1), MBP (ThermoFisher probe Mm01266402_m1), MyRF (Thermo Fisher probe Mm01194959_m1), and PDGFRA (ThermoFisher probe Mm00440701_m1). 100 ng of RNA per sample were input for cDNA synthesis using a Superscript IV cDNA synthesis kit (ThermoFisher 11750150) RT-PCR assays were run in 20 µL per reaction using SuperScript IV VILO master mix (ThermoFisher 11756050) on a QuantStudio 7 RT-PCR System (ThermoFisher). Marker gene enrichment at each time point was measured as fold change relative to expression at day 1 using the 2^-ΔΔCt^ method.^57^

#### Immunocytochemistry

O4 staining was performed on live cells prior to fixation. Cells were treated with anti-O4 antibody (R&D Systems, MAB1326) at a final concentration of 2 mg/mL in culture media for 45 – 90 minutes, 37°C, 5% CO_2_. Cells were gently washed, fixed at 4% paraformaldehyde for 15 minutes, and permeabilized/blocked in PBS containing 5% normal goat serum and 0.01 %Triton X-100 for a minimum of 60 minutes at room temperature. Antibody to human GPR17 (Millipore Sigma, HPA029766-100UL) was diluted 1:150 in permeabilization/block buffer (5% NGS, 0.01% Triton X-100 in PBS) and cells were stained with primary antibody overnight at +4°C. The following day, plates were washed and secondary antibodies were diluted 1:500 in PBS containing 1% NGS and 0.01% Triton X-100 and used at room temperature for 60 minutes with Alexa Fluor 488 anti-mouse IgM (Thermo Fisher, A-21042) and Alexa Fluor 647 anti-rabbit (Thermo Fisher, A-21245). Hoechst 33342 at 1:2000 was included in the secondary stain to identify nuclei.

#### Photolabeling and mass spectrometry analysis

PhotoClick cholesterol probe (700147P), 24(S)-hydroxycholesterol (700061P), and 24(R)-hydroxycholesterol (700071P) were obtained from Avanti. In gel-based photolabeling assay, GPR17 protein (WT construct, KC165 purified with SapA) was pre-treated with sterol as competitor for 1 hour at room temperature in 10 mM HEPES pH 7.5, 10 mM MgCl_2_, 500 mM KCl with gentle shaking, followed by incubation with 3 μM probe for 1 hour in the dark. Protein was then irradiated under 365 nm at 4 °C for 5 min three times in a Spectrolinker™ XL-1500 UV Crosslinker. After UV crosslinking, protein was conjugated to fluorophore via copper-catalyzed cyclo-addition (click chemistry) with 100 μM TAMRA-azide (tetramethylrhodamine azide, Lumiprobe B7130, CAS #825651-66-9), 10 mM CuSO_4_, 10 mM TCEP [Tris(2-carboxyethyl)phosphine hydrochloride, CAS#51805-45-9] and 100 μM TBTA (Tris[(1-benzyl-1H-1,2,3-triazol-4-yl)methyl]amine, CAS#510758-28-8) at room temperature 1h. NuPAGE™ LDS sample buffer was then added to each sample, separated by SDS-PAGE using Bis–Tris NuPAGE gels (4–12%, Invitrogen #NP0322), and MOPS running buffer (Life technologies #NP0002) in Xcell SureLock MiniCells (Invitrogen). Fluorescent labeling signals on gels were visualized by Typhoon and images were processed by ImageQuant. Protein loading in SDS–PAGE gels was visualized by silver stain.

For mass spectrometric analysis, 5μg GPR17 protein was pre-treated with 24S-OHC/ cholesterol for 1 hour, followed by incubation with 3 μM PhotoClick cholesterol probe at room temperature for 1 hour. After UV crosslinking, protein was denatured protein in 8 M urea, reduced with 10 mM DTT at 37 °C for 30 min and alkylated in 40 mM iodoacetamide. Urea was diluted to 1 M by adding 1 mM Tris pH 8.0 with 1 mM CaCl_2_. Protein was then digested by 0.5 μg trypsin overnight at 37 °C. Photo-labeled peptides were desalted on C18 stage tips, separated by EASY-nLC™ 1200 System with an EASY spray column (75 μm × 500 mm, 2 μm particle size, Thermo Scientific), and analyzed by Q Exactive HF Orbitrap LC-MS/MS. (Thermo Fisher Scientific). For peptide separation, a 10%-95% acetonitrile (ACN) gradient (solvent A: 0.1% FA/water, solvent B: 0.1% FA/ACN) was applied for 100 minutes as follow: 1– 60 minutes; 2%–30% solvent B, 1–75 minutes; 30%–45% solvent B, 75-85 minutes; 45%–98% solvent B, 85–87 minutes; isocratic elution at 98% solvent B, 87–91 minutes; 95%–2% solvent B, 91–92 minutes; 2% solvent B, 92–100 minutes. Standard full-MS/ dd-MS^2^ (Top10) method was applied for analysis: MS1 was acquired at resolution of 60,000 in the range of m/z = 300-2,200, AGC target 3E6. Top 10 ion precursors were selected for MS2 using data-dependent acquisition with charge exclusion of 1, ≥8. Fragmentation was performed with high-energy dissociation (HCD) with NCE: 27. Product ion spectra (MS2) were acquired at a resolution of 15,000, AGC 1E6, isolation window 1.6m/z. Data were searched against the sequence of human GPR17 using MaxQuant 2.1.3.0 software (Munich, Germany) with the following settings: MS/MS search tolerance of 20 ppm, 1% false discovery rate; maximum two missed cleavages, maximum five modifications per peptide; static modification of cysteine (+57.0215; iodoacetamide alkylation); variable modifications of methionine oxidation, N-terminal acetylation, and the adduct of cholesterol probe (+454.3447) at any amino acids. Detection of probe-modified peptides was further confirmed by manual examination for the presence of adducted fragment ions at corresponding retention time within 10 ppm mass accuracy in Xcalibur 2.2 (ThermoFisher). Photolabeling efficiency was estimated by generating selected ion chromatograms (SIC) of both unlabeled and photolabeled peptides, quantifying the area under curve and calculating efficiency as: labeled peptide / (unlabeled peptide + labeled peptide). Statistical significance was analyzed with Student’s paired t-test using GraphPad Prism 9.

#### Human brain sample sourcing

All human brain samples used in the study were obtained from the Edinburgh Brain Bank, centre of the UK Biobank. Details on informed consent and ethical oversight and approvals can be found at: https://www.ukbiobank.ac.uk/learn-more-about-uk-biobank/about-us/ethics. All sample were fresh frozen autopsy samples of cerebral white matter. Information on autopsy procedures, tissue processing, and banking can be found at: https://clinical-brain-sciences.ed.ac.uk/edinburgh-brain-bank.

#### GPR17 quantification by targeted proteomics

Frozen human brain tissues were obtained from Edinburgh Brain Bank (n=20 each group). Tissues were homogenized by MP bead homogenizer in 4% SDS, 0.1 M triethylammonium bicarbonate (TEAB) buffer with protease inhibitor cocktail (Roche, 5892791001). Brain homogenate was further sonicated by Covaris (two cycles of 250W, 10 seconds each). Lysate was centrifuged at 10,000 rpm for 10 minutes to remove insoluble debris. Protein concentration was measured by BCA assay and normalized to 2 mg/mL Lysate containing 20 μg of protein was reduced with 10 mM DTT at 37°C and then alkylated with 40 mM iodoacetamide for 60 min at 25°C with shaking. Samples were cleaned up by glass fiber material (Whatman Article No., 28418314) and protein was then digested in 0.1M TEAB by 1μg trypsin overnight at 37 °C. After digestion, five heavy isotope labeled, unique peptides of human GPR17 (customized order, Biosynth) were spiked into each sample at the ratio of 100 attomole heavy peptides per and 1 μg sample digests. Peptides were then desalted by Empore C18 membrane and separated by EASY-nLC™ 1200 System with an EASY spray column (75 μm × 500 mm, 2 μm particle size, Thermo Scientific) using a 140 min gradient as follow: 2%–30% solvent B, 1–105 minutes; 30%–45% solvent B, 105-121 minutes; 45%–98% solvent B, 121–126 minutes; isocratic elution at 98% solvent B, 126–131 minutes; 95%–2% solvent B, 131–132 minutes; 2% solvent B, 132–140 minutes. Samples were analyzed by Q Exactive HF Orbitrap LC-MS/MS (Thermo Fisher Scientific) using parallel-reaction monitoring (PRM) mode with the following settings: MS2 resolution of 30,000, AGC target 5E5, isolation window 1.0 m/z, maximum IT 200ms, scheduled acquisition window and NCE optimized for each targeted peptide. PRM data were analyzed in Skyline software and light-to-heavy (L/H) ratios were obtained. Protein concentration was calculated from the averaged L/H ratio of target peptides (HALCNLLCGK, ILALANR, FLAIVHPVK, SVYVLHYR, TNESSLSAK) and protein extraction efficiency in each tissue sample.

#### Quantification of 24S-hydroxycholesterol

The lipids reference standards were purchased from Avanti Research (Birmingham, AL, US) and GC derivatization reagent, N, O-Bis (trimethylsilyl) trifluoroacetamide with trimethylchlorosilane was acquired from Millipore Sigma (Burlington, MA, US) Brain tissue samples were homogenized in methanol (25 mg tissue per mL methanol) using a FastPrep system (MP Biomedicals, CA). For lipid extraction, 0.08 mL of the homogenate, equivalent to 2 mg of tissue, was utilized. The lipid extraction process was carried out using the Folch method ^58^, which employed a biphasic solvent system consisting of chloroform, methanol, and water. Briefly, 0.2 mL of chloroform and 0.03 mL of water were added to 0.08 mL tissue homogenate. Additionally, 0.01 mL of the surrogate, deuterated MAS-412-d7 (0.0025 mg/mL methanol), was spiked into each sample to monitor lipid extraction recovery. The samples were agitated in a Thermomixer at room temperature for approximately 20-30 min, followed by centrifugation to divide the two liquid phases. The lipid-rich chloroform phase at the bottom was carefully transferred into a clean glass vial. Subsequently, 0.2 mL of chloroform was added to the residual sample for a second extraction. The lipid-rich phases from both extractions were then combined and evaporated under a stream of nitrogen gas. The resulting lipid extracts were reconstituted in 0.02 mL of the derivatization reagent, N, O-Bis (trimethylsilyl) trifluoroacetamide with trimethylchlorosilane, and incubated at 60°C for 20 minutes. Before GC-MS analysis, internal standards, including zymostenol-d7, lathosterol-d7, and MAS-414-d6, were added to each sample.

A 1 µL volume was analyzed using an Agilent 7890A gas chromatograph coupled with a 5975C inert MSD single quadrupole mass spectrometer, equipped with an Electron Impact (EI) ion source. The 24S-hydroxycholesterol (24S-OHC) was separated from other hydroxycholesterol isomers, such as 7α-OHC, 7ß-OHC, 4ß-OHC, 20α-OHC, 22S-OHC, 22R-OHC, 25-OHC, and 27-OHC, using an HP-5MS capillary column (60 m × 0.25 mm × 0.25 µm). Note: the enantiomer pair, 24S-OHC and 24R-OHC, could not be separated using this GC column. The 24-OHC measurement was labeled as 24S-OHC because 24S-OHC is the predominant cholesterol metabolite in the brain, and no literature brain data on 24R-OHC is available. GC-MS data were collected in Selected Ion Monitoring (SIM) scan mode with positive polarity and processed using Agilent MassHunt (version B.07.01. SP). The GC-MS peak area for 24S-OHC was integrated and its ratio to the internal standard (IS) was calculated. These ratios were then converted to nanomoles per gram of tissue based on the analyte’s calibration curve. The calibration range of 24S-OHC spanned from 20 ng/mL to 20 µg/mL.

## Acknowledgments

The authors thank Professor Cheng Zhang for productive conversations on protein production for structure.

## References

1. Chen, Y. et al. The oligodendrocyte-specific G protein-coupled receptor GPR17 is a cell-intrinsic timer of myelination. Nat Neurosci 12, 1398–406 (2009).

2. Merten, N., et al. Repurposing HAMI3379 to Block GPR17 and Promote Rodent and Human Oligodendrocyte Differentiation. Cell Chem Biol 25, 775–786.e5 (2018).

3. Simon, K. et al. The Orphan G Protein-coupled Receptor GPR17 Negatively Regulates Oligodendrocyte Differentiation via G&#x3b1;i/o and Its Downstream Effector Molecules *. Journal of Biological Chemistry 291, 705–718 (2016).

4. Raffaele, S. et al. Characterisation of GPR17-expressing oligodendrocyte precursors in human ischaemic lesions and correlation with reactive glial responses. J Pathol 265, 226–243 (2025).

5. Jakel, S. et al. Altered human oligodendrocyte heterogeneity in multiple sclerosis. Nature 566, 543–547 (2019).

6. Macnair, W. et al. snRNA-seq stratifies multiple sclerosis patients into distinct white matter glial responses. Neuron 113, 396–410 e9 (2025).

7. Häberlein, F. et al. Humanized zebrafish as a tractable tool for in vivo evaluation of pro-myelinating drugs. Cell Chem Biol 29, 1541–1555.e7 (2022).

8. Lu, C. et al. G-Protein-Coupled Receptor Gpr17 Regulates Oligodendrocyte Differentiation in Response to Lysolecithin-Induced Demyelination. Sci Rep 8, 4502 (2018).

9. Marucci, G. et al. GPR17 receptor modulators and their therapeutic implications: review of recent patents. Expert Opin Ther Pat 29, 85–95 (2019).

10. Martin, A.L., Steurer, M.A. & Aronstam, R.S. Constitutive Activity among Orphan Class-A G Protein Coupled Receptors. PLOS ONE 10, e0138463 (2015).

11. Hennen, S. et al. Decoding Signaling and Function of the Orphan G Protein–Coupled Receptor GPR17 with a Small-Molecule Agonist. Science Signaling 6, ra93–ra93 (2013).

12. Qi, A.D., Harden, T.K. & Nicholas, R.A. Is GPR17 a P2Y/leukotriene receptor? examination of uracil nucleotides, nucleotide sugars, and cysteinyl leukotrienes as agonists of GPR17. J Pharmacol Exp Ther 347, 38–46 (2013).

13. Harrington, A.W. et al. Identification and characterization of select oxysterols as ligands for GPR17. British Journal of Pharmacology 180, 401–421 (2023).

14. Lin, X. et al. Structural basis of ligand recognition and self-activation of orphan GPR52. Nature 579, 152–157 (2020).

15. Liu, G., Li, X., Wang, Y., Zhang, X. & Gong, W. Structural basis for ligand recognition and signaling of the lysophosphatidylserine receptors GPR34 and GPR174. PLOS Biology 21, e3002387 (2023).

16. Ye, F., et al. Cryo-EM structure of G-protein-coupled receptor GPR17 in complex with inhibitory G protein. MedComm 3, e159 (2022).

17. Christa E. Mueller, C.P., Michael Louis Robert Deligny, Ali El-Tayeb, Joerg Hockemeyer, Marie Ledecq, Joël Mercier, Laurent Provins, Nader M. Boshta, Sanjay Bhattarai, Vigneshwaran Namasivayam, Mario Funke, Lukas Schwach, Sabrina Gollos, Daniel Von Laufenberg, Anaïs Barré. Aza (indole)-, benzothiophene-, and benzofuran-3-sulfonamides. Vol. 2 (ed. Organization, W.I.P.) (UCB PHARMA GMBH, International, 2018).

18. Baqi, Y. et al. 3-(2-Carboxyethyl)indole-2-carboxylic Acid Derivatives: Structural Requirements and Properties of Potent Agonists of the Orphan G Protein-Coupled Receptor GPR17. Journal of Medicinal Chemistry 61, 8136–8154 (2018).

19. Zhang, K. et al. Structure of the human P2Y12 receptor in complex with an antithrombotic drug. Nature 509, 115–118 (2014).

20. Gusach, A. et al. Structural basis of ligand selectivity and disease mutations in cysteinyl leukotriene receptors. Nat Commun 10, 5573 (2019).

21. Luginina, A. et al. Structure-based mechanism of cysteinyl leukotriene receptor inhibition by antiasthmatic drugs. Sci Adv 5, eaax2518 (2019).

22. Chen, H., Huang, W. & Li, X. Structures of oxysterol sensor EBI2/GPR183, a key regulator of the immune response. Structure 30, 1016–1024.e5 (2022).

23. Chun, E. et al. Fusion Partner Toolchest for the Stabilization and Crystallization of G Protein-Coupled Receptors. Structure 20, 967–976 (2012).

24. Zhang, K., Wu, H., Hoppe, N., Manglik, A. & Cheng, Y. Fusion protein strategies for cryo-EM study of G protein-coupled receptors. Nature Communications 13, 4366 (2022).

25. Gusach, A. et al. Structural basis of ligand selectivity and disease mutations in cysteinyl leukotriene receptors. Nature Communications 10, 5573 (2019).

26. Mukherjee, S. et al. Synthetic antibodies against BRIL as universal fiducial marks for single−particle cryoEM structure determination of membrane proteins. Nature Communications 11, 1598 (2020).

27. Zhu, H. et al. Discovery of novel and selective GPR17 antagonists as pharmacological tools for developing new therapeutic strategies in diabetes and obesity. Eur J Med Chem 295, 117794 (2025).

28. Boshta, N.M. et al. Discovery of Anthranilic Acid Derivatives as Antagonists of the Pro-Inflammatory Orphan G Protein-Coupled Receptor GPR17. Journal of Medicinal Chemistry 67, 19365–19394 (2024).

29. White, K.L. et al. Structural Connection between Activation Microswitch and Allosteric Sodium Site in GPCR Signaling. Structure 26, 259–269.e5 (2018).

30. Wu, Z. et al. Dynamic Insights into the Self-Activation Pathway and Allosteric Regulation of the Orphan G-Protein-Coupled Receptor GPR52. Journal of Chemical Information and Modeling 63, 5847–5862 (2023).

31. Hoppe, N. et al. GPR161 structure uncovers the redundant role of sterol-regulated ciliary cAMP signaling in the Hedgehog pathway. Nature Structural & Molecular Biology 31, 667–677 (2024).

32. Fan, Y., et al. Allosteric coupling between G-protein binding and extracellular ligand binding sites in GPR52 revealed by 19F-NMR and cryo-electron microscopy. MedComm 4, e260 (2023).

33. Colquhoun, D. Why the Schild method is better than Schild realised. Trends in Pharmacological Sciences 28, 608–614 (2007).

34. Syed, Y.A. et al. Inhibition of phosphodiesterase-4 promotes oligodendrocyte precursor cell differentiation and enhances CNS remyelination. EMBO Mol Med 5, 1918–34 (2013).

35. Hulce, J.J., Cognetta, A.B., Niphakis, M.J., Tully, S.E. & Cravatt, B.F. Proteome-wide mapping of cholesterol-interacting proteins in mammalian cells. Nat Methods 10, 259–64 (2013).

36. Lum, K.M. et al. Mapping Protein Targets of Bioactive Small Molecules Using Lipid-Based Chemical Proteomics. ACS Chem Biol 12, 2671–2681 (2017).

37. Budelier, M.M. et al. Photoaffinity labeling with cholesterol analogues precisely maps a cholesterol-binding site in voltage-dependent anion channel-1. Journal of Biological Chemistry 292, 9294–9304 (2017).

38. Hill, R.A., Li, A.M. & Grutzendler, J. Lifelong cortical myelin plasticity and age-related degeneration in the live mammalian brain. Nat Neurosci 21, 683–695 (2018).

39. Hughes, E.G., Orthmann-Murphy, J.L., Langseth, A.J. & Bergles, D.E. Myelin remodeling through experience-dependent oligodendrogenesis in the adult somatosensory cortex. Nat Neurosci 21, 696–706 (2018).

40. Yeung, M.S. et al. Dynamics of oligodendrocyte generation and myelination in the human brain. Cell 159, 766–74 (2014).

41. Lees, J.A. et al. An inverse agonist of orphan receptor GPR61 acts by a G protein-competitive allosteric mechanism. Nature Communications 14, 5938 (2023).

42. Frauenfeld, J. et al. A saposin-lipoprotein nanoparticle system for membrane proteins. Nature Methods 13, 345–351 (2016).

43. Lloris-Garcerá, P. et al. DirectMX – One-Step Reconstitution of Membrane Proteins From Crude Cell Membranes Into Salipro Nanoparticles. Frontiers in Bioengineering and Biotechnology 8(2020).

44. Zhuang, Y. et al. Structure of formylpeptide receptor 2-Gi complex reveals insights into ligand recognition and signaling. Nature Communications 11, 885 (2020).

45. Liu, P. et al. The structural basis of the dominant negative phenotype of the Gαi1β1γ2 G203A/A326S heterotrimer. Acta Pharmacologica Sinica 37, 1259–1272 (2016).

46. Mukherjee, S. et al. Synthetic antibodies against BRIL as universal fiducial marks for single-particle cryoEM structure determination of membrane proteins. Nat Commun 11, 1598 (2020).

47. Ereno-Orbea, J. et al. Structural Basis of Enhanced Crystallizability Induced by a Molecular Chaperone for Antibody Antigen-Binding Fragments. J Mol Biol 430, 322–336 (2018).

48. Tsutsumi, N. et al. Structure of human Frizzled5 by fiducial-assisted cryo-EM supports a heterodimeric mechanism of canonical Wnt signaling. Elife 9(2020).

49. Punjani, A., Rubinstein, J.L., Fleet, D.J. & Brubaker, M.A. cryoSPARC: algorithms for rapid unsupervised cryo-EM structure determination. Nat Methods 14, 290–296 (2017).

50. Rubinstein, J.L. & Brubaker, M.A. Alignment of cryo-EM movies of individual particles by optimization of image translations. J Struct Biol 192, 188–95 (2015).

51. Punjani, A., Zhang, H. & Fleet, D.J. Non-uniform refinement: adaptive regularization improves single-particle cryo-EM reconstruction. Nat Methods 17, 1214–1221 (2020).

52. Liebschner, D. et al. Macromolecular structure determination using X-rays, neutrons and electrons: recent developments in Phenix. Acta Crystallogr D Struct Biol 75, 861–877 (2019).

53. Krishna Kumar, K., et al. Structure of a Signaling Cannabinoid Receptor 1-G Protein Complex. Cell 176, 448–458 e12 (2019).

54. Pettersen, E.F. et al. UCSF Chimera--a visualization system for exploratory research and analysis. J Comput Chem 25, 1605–12 (2004).

55. Emsley, P., Lohkamp, B., Scott, W.G. & Cowtan, K. Features and development of Coot. Acta Crystallogr D Biol Crystallogr 66, 486–501 (2010).

56. Arunlakshana, O. & Schild, H.O. Some quantitative uses of drug antagonists. Br J Pharmacol Chemother 14, 48–58 (1959).

57. Livak, K.J. & Schmittgen, T.D. Analysis of Relative Gene Expression Data Using Real-Time Quantitative PCR and the 2−ΔΔCT Method. Methods 25, 402–408 (2001).

58. Folch, J., Lees, M. & Sloane Stanley, G.H. A simple method for the isolation and purification of total lipides from animal tissues. J Biol Chem 226, 497–509 (1957).

