## Extended Data Figure Legends for "GPR17 structure and agonism with small molecules and oxysterols"

**Extended Data Figure 1.** Biochemical preparation of GPR17. a) Agonist complex in LMNG detergent, representative size exclusion profile (left) and SDS PAGE gel (right). b) Antagonist complex in SapA nanodisc, representative size exclusion profile (left) and SDS PAGE gel (right).

**Extended Data Figure 2.** CryoEM Workflow and Data Quality for MDL- and I-185-bound GPR17 structures. a, b) CryoEM data processing workflow for MDL- and I-185-bound structures, respectively. c, d) CryoEM maps of MDL- and I-185-bound structures, respectively, colored by relative local resolution according to scale. e, f) CryoEM density for MDL- and I-185-bound structures, respectively, showing each of the 7TM helices and H8.

**Extended Data Figure 3**. Antagonist binding site in d-GPCRs. GPR17 +I-185 in magenta, CysLT1 +pranlukast (PDB 6rz4) in deep purple, CysLT2 +ONO-2570366 (PDB 6rz6) in orange. a) Comparing ligand placement, view from EC, with ECL2 highlighted, showing side occupation of the pocket only by I-185. b) Showing GPR17 ECL2 depth location with respect to pocket and ligand, circle highlights ECL2 clash with pranlukast (deep purple) and ONO-2570366 (orange). c) Surface view showing groove between TM4 and TM5 into which antagonists extend. d) Cartoon cylinder depiction to show an outward shift in TM5 of GPR17 due to I-I85 compared to shift inwards towards antagonists of CysLTs, and depth of GPR17 ECL2 pocket, despite close alignment with CysLTs on beta turn.

**Extended Data Figure 4**. GPR17 and EBI2. GPR17 + MDL in forest, EBI2 (GPR183) + CHS (PDB 7tuz) in sky blue. a) GPR17 surface view showing groove between TM4 and TM5 that is the proposed access point for hydrophobic substrates and that agonists extend into. b) Comparing ligand placement (view from EC with surface transparency) showing side occupation of the pocket only by MDL. c) Electrostatic surface representation highlights nonpolar groove of MDL-bound GPR17 (left, green MDL) and 24S-bound EBI2 (right), the putative sterol entrance for EBI2. d) Cartoon cylinder depiction to show an outward shift in TM6 of EBI2 compared to that of GPR17 and changes in ECL2 placement.

**Extended Data Figure 5**. Comparison of P2Y_12_ and GPR17 global conformational changes upon activation. a) P2Y_12_ active state (sage, PDB 7xxi) and inactive state (salmon, PDB 4ntj). b) GPR17 active state (green) and inactive state (fuchsia).

**Extended Data Figure 6**. Comparison of self-activated GPCRs, ligand binding sites, ECL2 changes and surface coverage. a) Ligands shown I-185-bound GPR17 (white cartoon): I-185 (magenta), GSK682753A (EBI2, PDB 7tuy, sky blue), C17 (GPR52, PDB 8hmp, orange), CLR (GPR161, PDB 8kh4, dark purple), CLR (GPR161, PDB 8smv, light purple), compound1/ZOB (GPR61, PDB 8tb7, lime green). b) ECL2 comparisons for GPR17 (magenta, dot and cartoon representation) and EBI2 +GSK682753A (PDB 7tuy, sky blue, left), GPR52 +C17 (PDB 8hmp, orange, center), and GPR161 (PDB 8smv, light purple, right). TM6 and TM7 are transparent in the side view (top row) for visibility of ECL2 depth in orthosteric pocket.

**Extended data Figure 7.** Characterization of GPR17 in human OPC culture. a-d) qPCR data showing enrichment of GPR17; PDGFRA, a progenitor marker; MyRF, an early and late differentiation marker; and MBP, a myelin protein marker over the course of one week of culture. Note the enrichment of GPR17 at 5 days in culture. e) Immunohistochemistry images of the early oligodendrocyte differentiation marker O4 with the nuclei stain DAPi demonstrating near uniformity of oligodendrocytes in the preoligodendrocyte stage at 5 days in culture. f) Immunohistochemical staining of GPR17 demonstrating that nearly all cells are GPR17 positive at 5 days in culture. Plots and error bars in a to d are mean ± SEM. N=3 independent experiments. Scale bar = 100 µm.

**Extended data Figure 8.** Generation of GPR17 KO approach. a) Sequence and location of guide RNAs and genotyping primers for GPR17 exon 2 deletion region. b) Reduction of GPR17 gene expression confirmed by 2 separate Taqman qPCR probes. c) Representative images of IHC staining with DAPI nuclei dye demonstrating absence of GPR17 protein expression in brain in GPR17 KO versus WT brain. d) Double IHC staining for GPR17 (red) and the pan oligodendrocyte marker gene OLIG2 (green) confirming GPR17 deletion and absence from oligodendrocyte cells in the GPR17 KO mice. Plots and error bars in b are mean ± SEM; N=4 mice. Scale bar = 100 µm.
