## Extended Data Figures for "GPR17 structure and agonism with small molecules and oxysterols"

### Slide 1
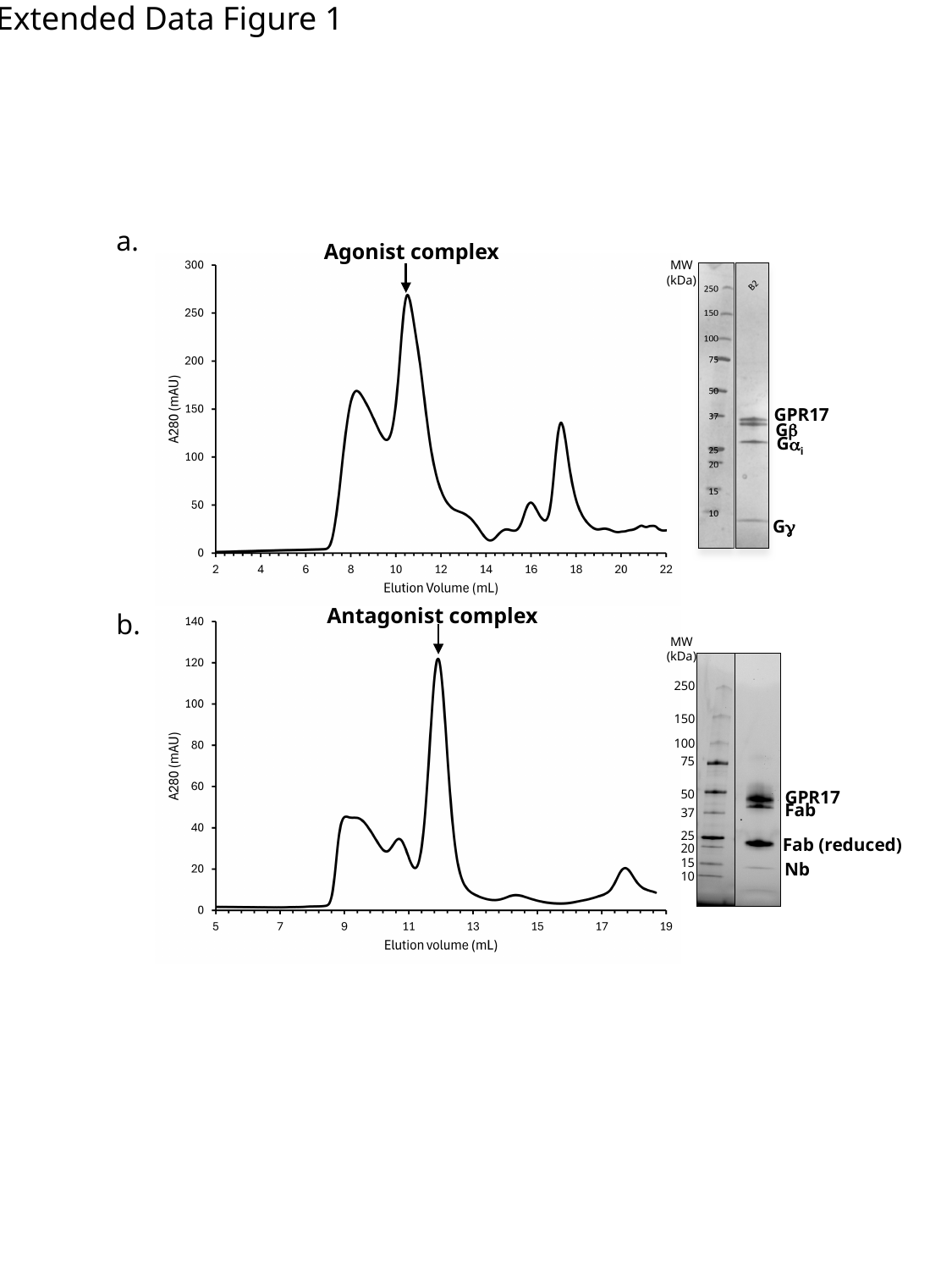

Extended Data Figure 1
a.
Agonist complex
MW
(kDa)
GPR17
Gβ
Gα_i_
Gγ
Antagonist complex
b.
MW
(kDa)
250
150
100
75
GPR17
50
Fab
37
25
Fab (reduced)
20
15
Nb
10

### Slide 2
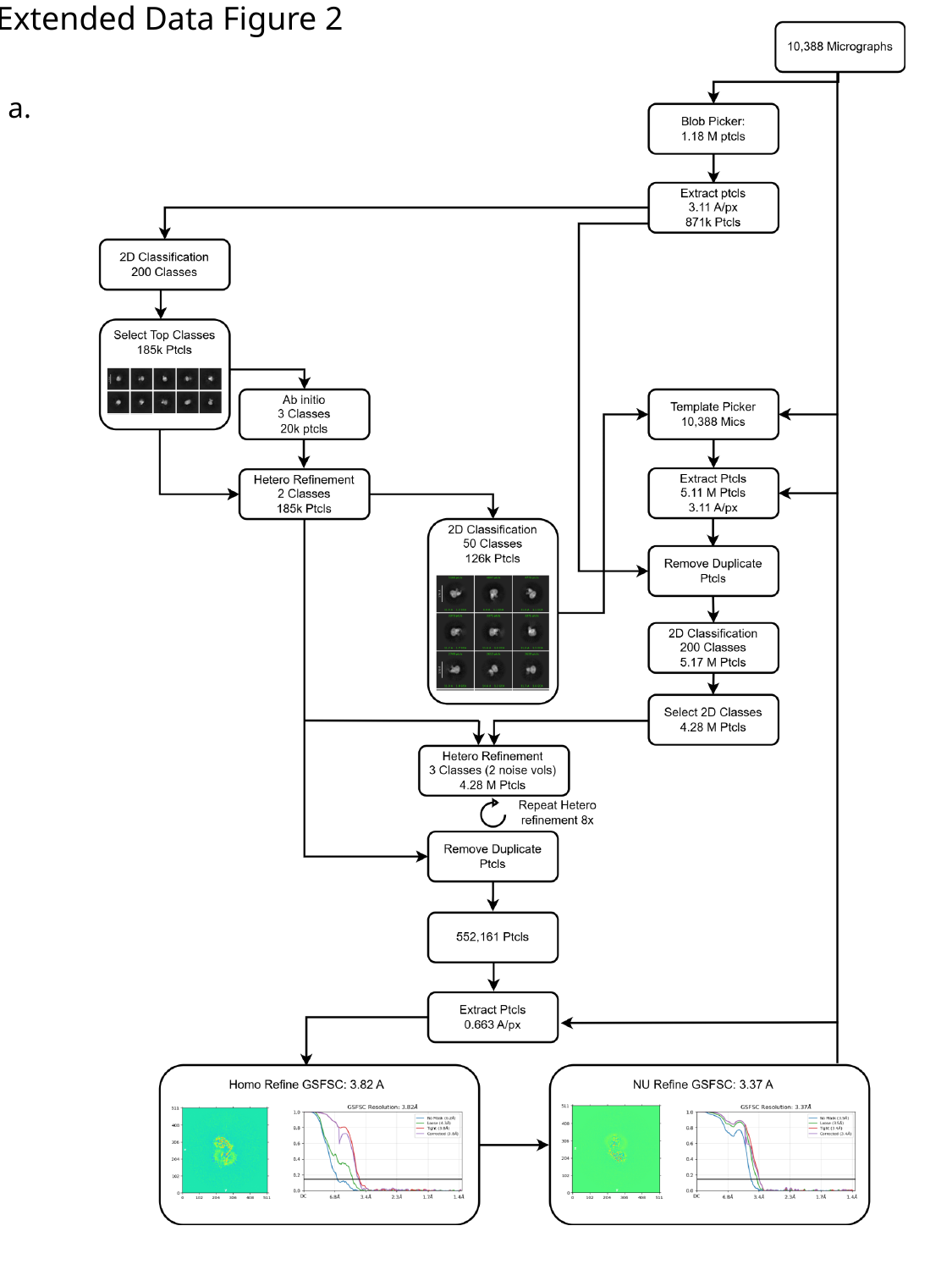

Extended Data Figure 2
a.

### Slide 3
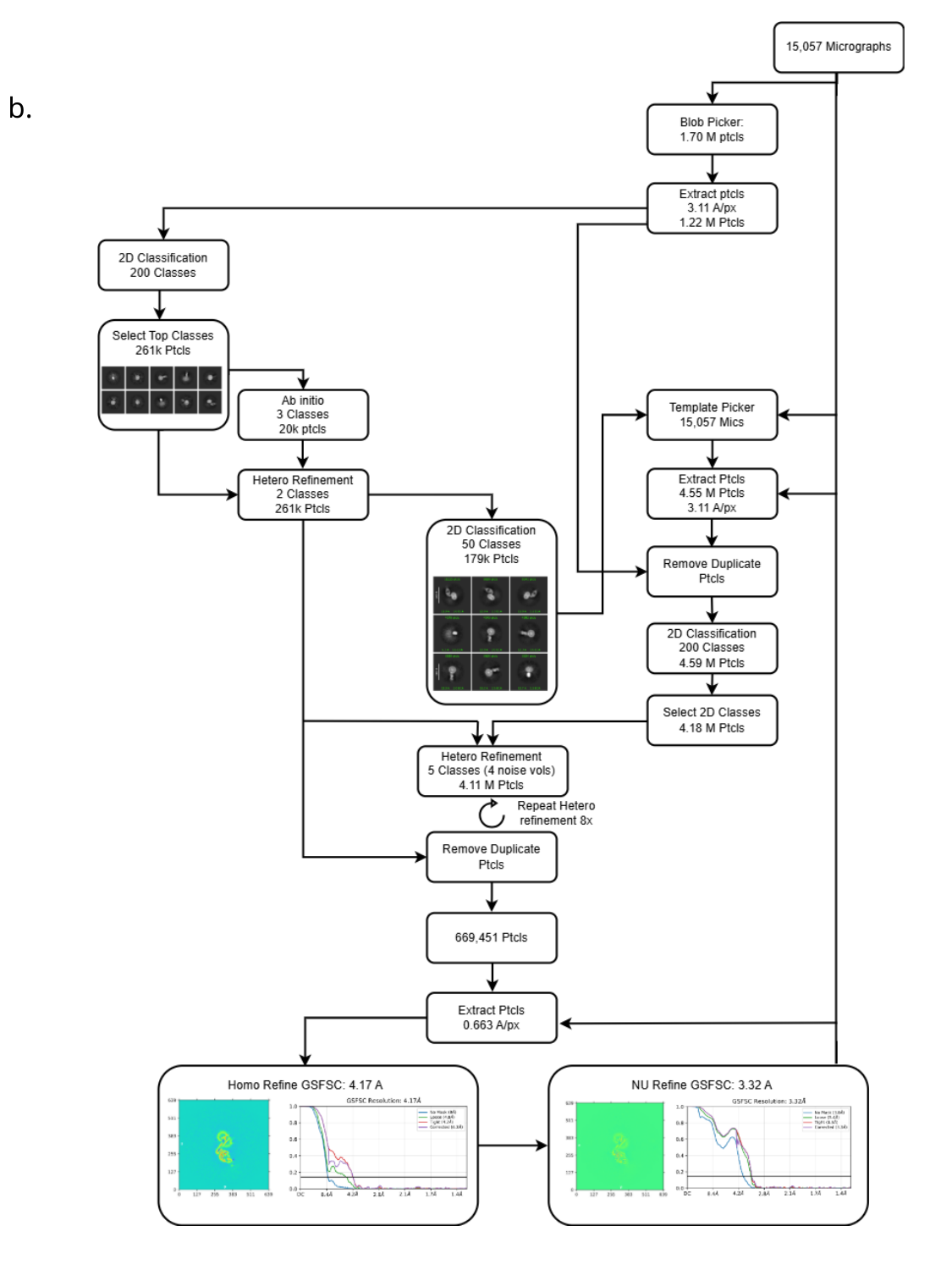

b.

### Slide 4
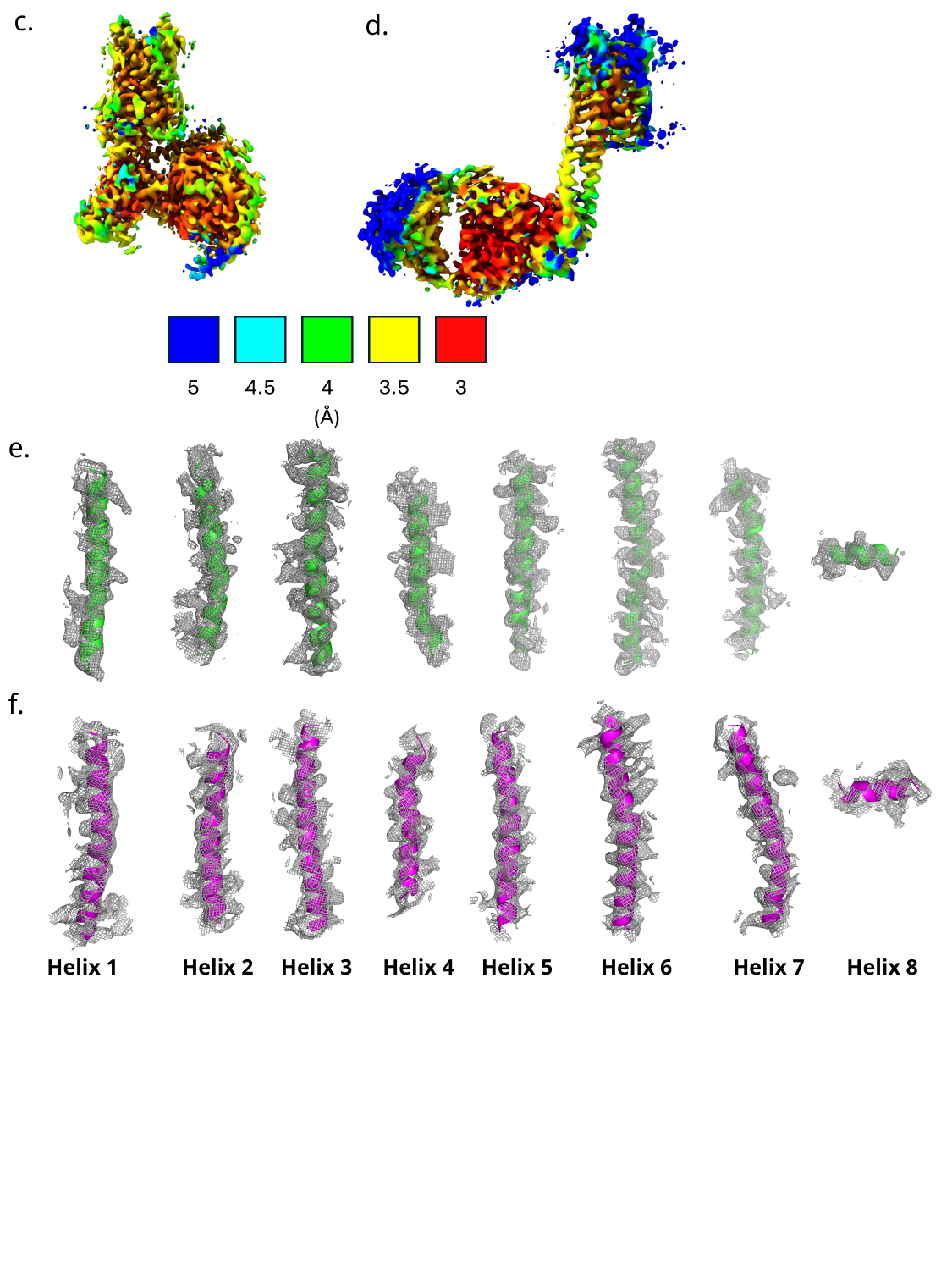

c.
d.
e.
f.
Helix 1
Helix 2
Helix 3
Helix 4
Helix 5
Helix 6
Helix 7
Helix 8

### Slide 5
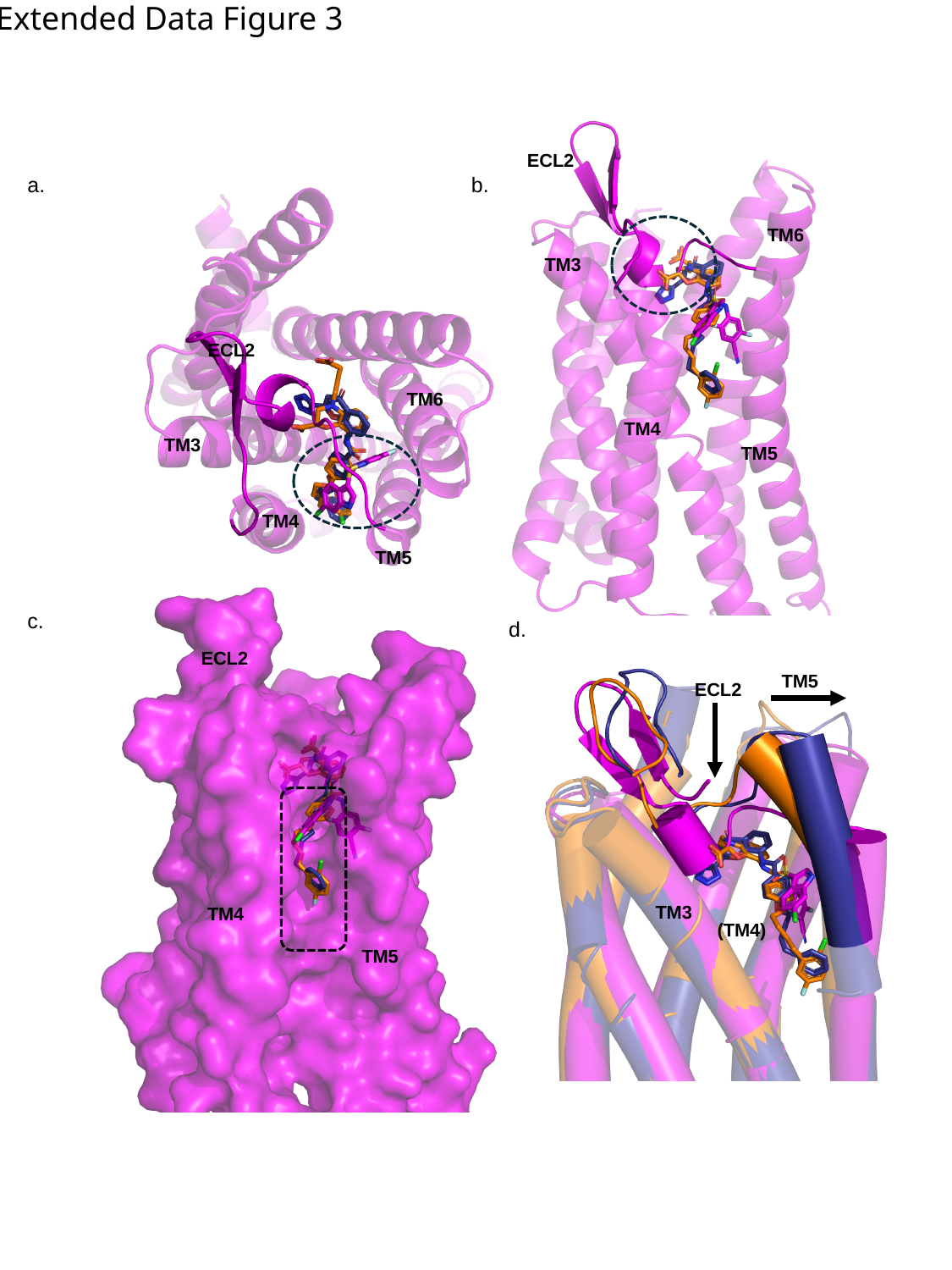

Extended Data Figure 3
ECL2
b.
a.
TM6
TM3
ECL2
TM6
TM4
TM3
TM5
TM4
TM5
c.
d.
ECL2
TM5
ECL2
TM3
TM4
(TM4)
TM5

### Slide 6
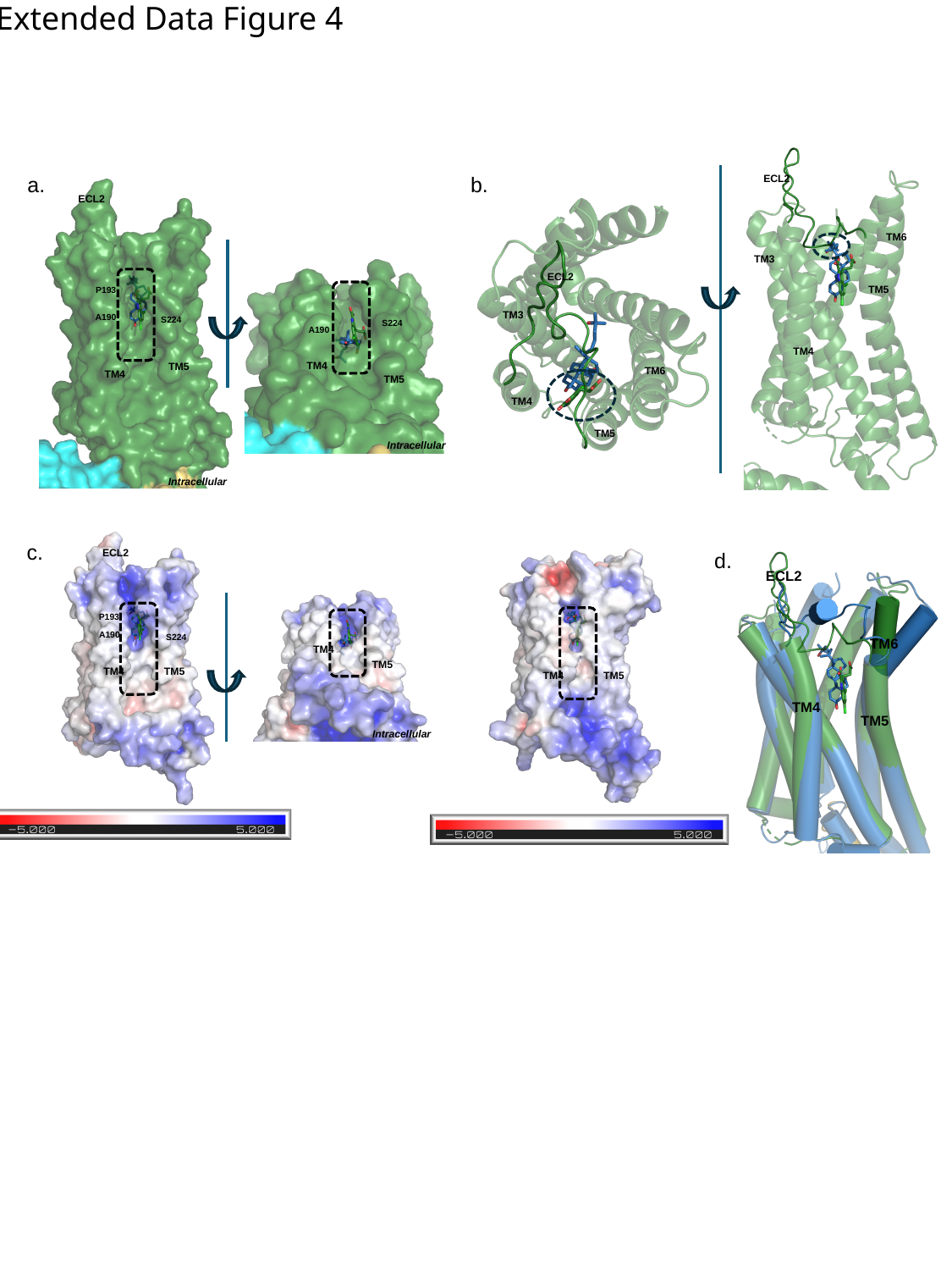

Extended Data Figure 4
b.
a.
ECL2
ECL2
TM6
TM3
ECL2
TM5
P193
TM3
A190
S224
S224
A190
TM4
TM4
TM5
TM6
TM4
TM5
TM4
TM5
Intracellular
Intracellular
c.
ECL2
d.
ECL2
P193
A190
S224
TM6
TM4
TM5
TM4
TM5
TM4
TM5
TM4
TM5
Intracellular

### Slide 7
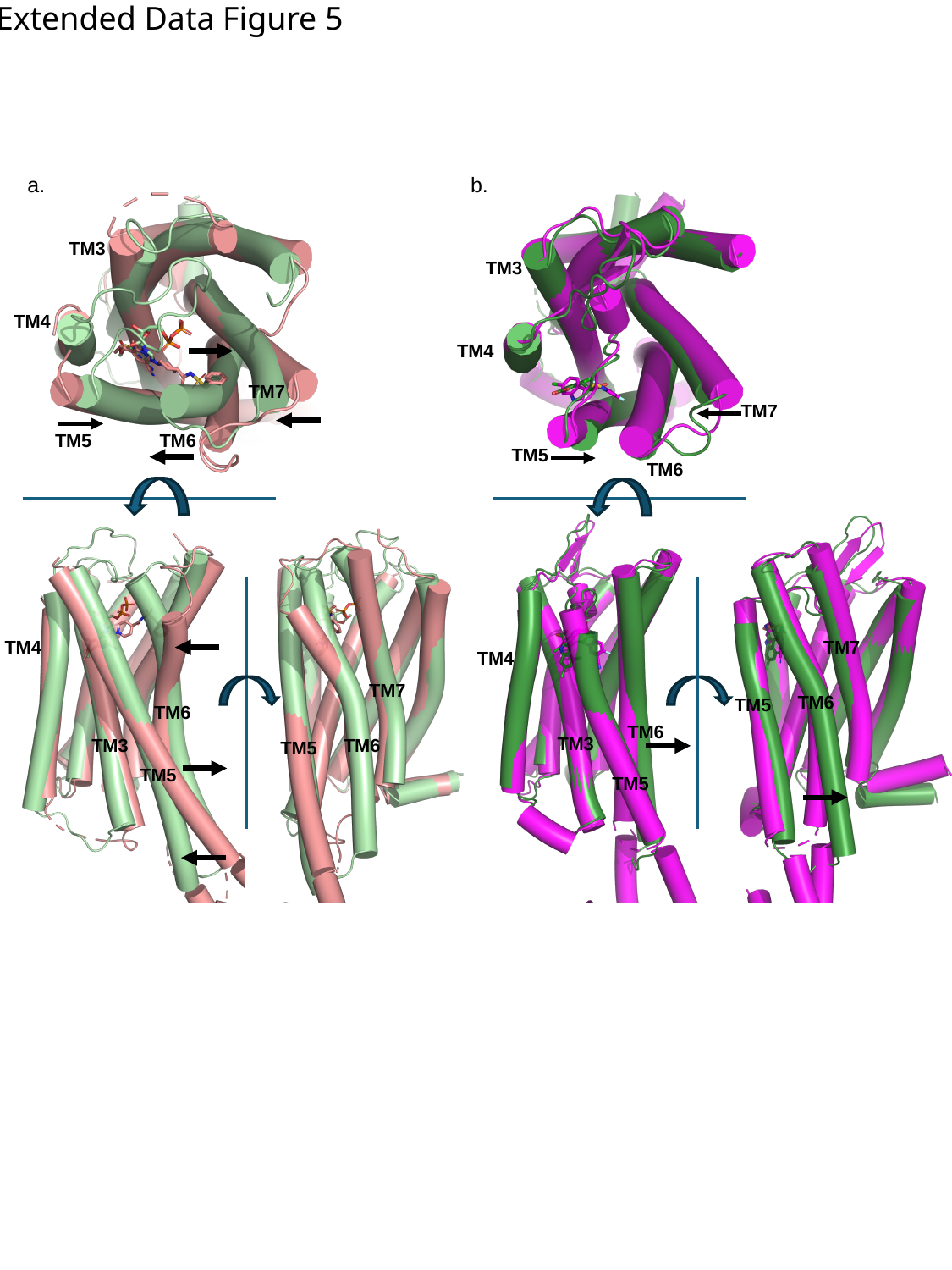

Extended Data Figure 5
b.
a.
TM3
TM3
TM4
TM4
TM7
TM7
TM5
TM6
TM5
TM6
TM4
TM7
TM4
TM7
TM6
TM5
TM6
TM6
TM3
TM3
TM6
TM5
TM5
TM5

### Slide 8
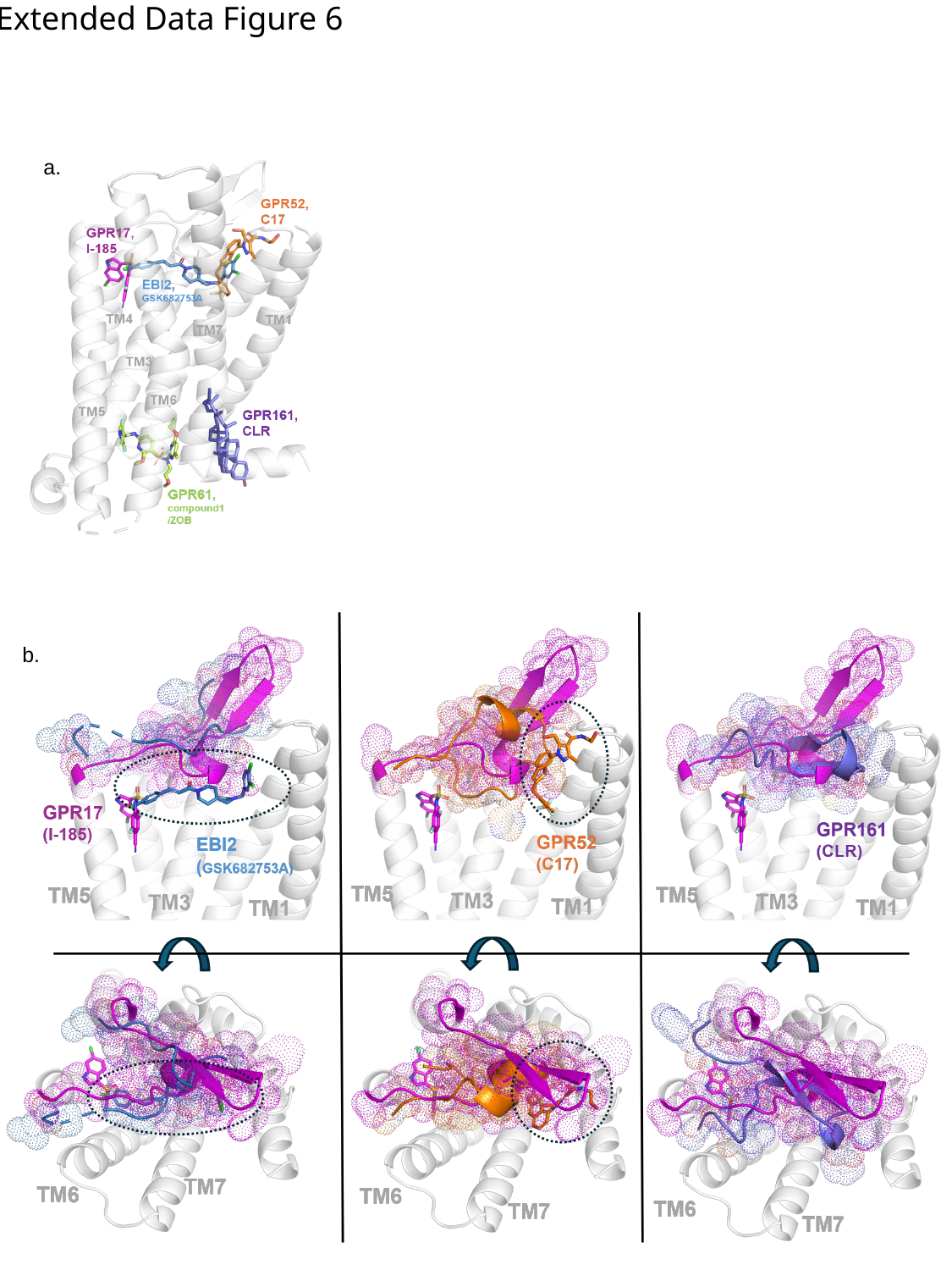

Extended Data Figure 6
a.
b.

### Slide 9
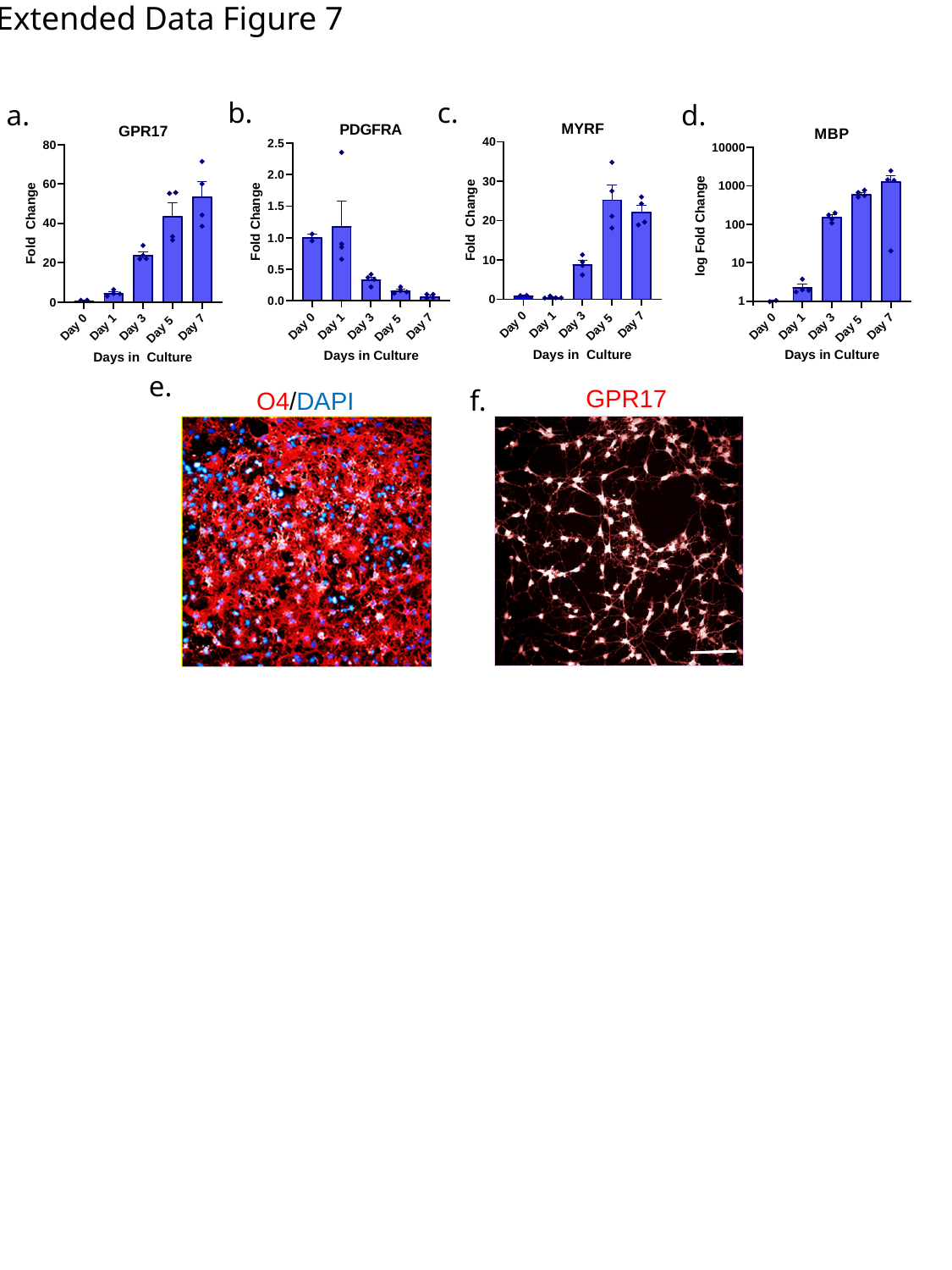

Extended Data Figure 7
b.
c.
a.
d.
e.
f.
GPR17
O4/DAPI

### Slide 10
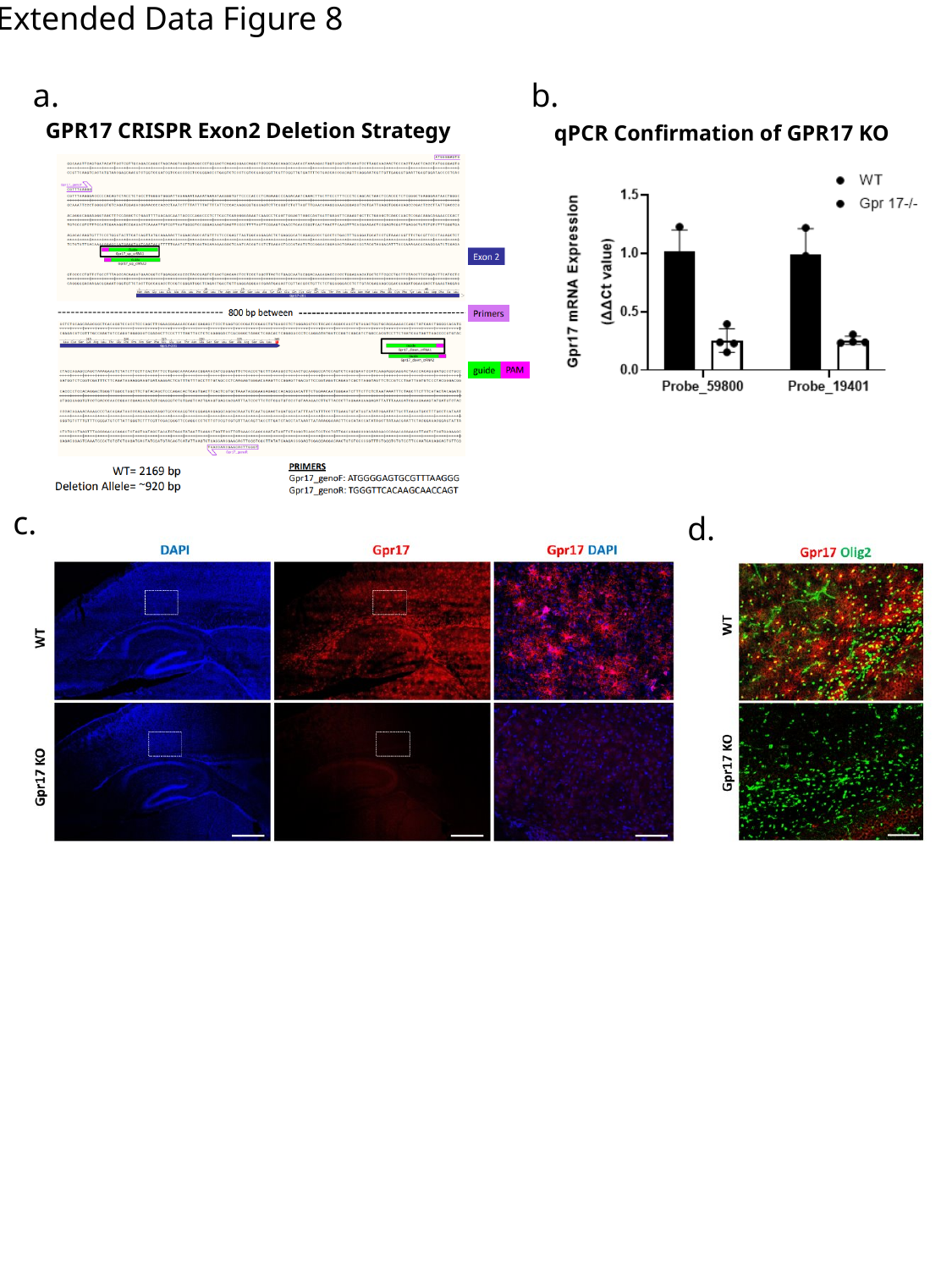

Extended Data Figure 8
a.
b.
GPR17 CRISPR Exon2 Deletion Strategy
qPCR Confirmation of GPR17 KO
c.
d.
